# BASELINE: A CRISPR Base Editing Platform for Mammalian-Scale Single-Cell Lineage Tracing

**DOI:** 10.1101/2025.03.19.644238

**Authors:** Evan Winter, Francesco Emiliani, Aidan Cook, Asma Abderrahim, Aaron McKenna

## Abstract

A cell’s fate is shaped by its inherited state, or lineage, and the ever-shifting context of its environment. CRISPR-based recording technologies are a promising solution to map the lineage of a developing system, yet challenges remain regarding single-cell recovery, engineering complexity, and scale. Here, we introduce BASELINE, which uses base editing to generate high-resolution lineage trees in conjunction with single-cell profiling. BASELINE uses the Cas12a adenine base editor to irreversibly edit nucleotides within 50 synthetic target sites, which are integrated multiple times into a cell’s genome. We show that BASELINE accumulates lineage-specific marks over a wide range of biologically relevant intervals, recording more than 4300 bits of information in a model of pancreatic cancer, a 50X increase over existing technologies. Single-cell sequencing reveals high-fidelity capture of these recorders, recovering lineage reconstructions up to 40 cell divisions deep, within the estimated range of mammalian development. We expect BASELINE to apply to a wide range of lineage-tracing projects in development and disease, especially in which cellular engineering makes small, more distributed systems challenging.

## Introduction

A shared lineage links each cell within a multicellular organism. Efforts to trace lineages date back to the 19th century^1,2^ when scientists discovered that cells divide from pre-existing cells. Improvements in microscopy^3^, the advent of fluorescent reporters^4,5^, and DNA sequencing technologies^6,7^ have improved phylogenetic cellular reconstruction. Recent advances in lineage tracing have leveraged DNA nucleases such as CRISPR/Cas9 to generate dsDNA breaks within a compact array of synthetic target sites. The dsDNA breaks are repaired and cleaved repeatedly until the DNA repair process induces errors in the form of insertions or deletions (indels), which are inherited by daughter cells. First described in the GESTALT^8^ and MEMOIR^9^ papers, this approach has been used to trace lineage in various biological contexts^10–13^. Unfortunately, these systems have drawbacks. If multiple targets within an array are cleaved simultaneously, the DNA between the cleavages, including any previous editing information, is lost. Additionally, these nucleases can activate a p53 response^14^, changing the underlying cellular phenotype^15^.

Researchers have turned to alternative genome editing approaches to overcome these limitations and enable more precise phylogenetic reconstructions. Base editors^16,17^ and prime editors^18^, originally developed for targeted DNA modifications without double-stranded breaks, have since been adapted for CRISPR/Cas-based lineage tracing^19–21^. In addition to these Cas-based tools, integrases^22^ and homing endonucleases^23^ are viable and capable of lineage recording. While the promise of these technologies remains high, there are significant challenges. Prime editors still have editing efficiency issues^24^, limiting applicability in many cell types and leading to lower-resolution phylogenetic trees. Additionally, current homing endonuclease systems cannot be captured simultaneously with gene expression data, making it challenging to pair cell phenotype and trajectory with lineage information^23^. Lastly, integrases still create multi-site deletions^22^ and recorder recovery varies drastically between technologies^25^. There is a need for a lineage tracing technology with low cellular engineering requirements, high recorder diversity, high recorder recovery rate, and high information density.

Here, we introduce BASELINE, a <u>base</u> editing <u>line</u>age tracing system that uses Cas12a adenosine base editors to create predictable and irreversible changes to lineage recorders (**Fig. 1A**). Our recorder design utilizes 50 unique and synthetic target sites, which in total contain 272 adenosines (2^272^ or >7.58*10^81^ outcomes). With Cas12a, we can multiplex the RNA guides, enabling the compact packaging of the guide arrays and recorder onto a single construct (**Fig. 1B**). High recorder expression is coupled with T7 in vitro transcription (IVT) to maximize recorder recovery without PCR bias^26^. To sequence our 1.5 kbp recorders, we use Oxford Nanopore Technology (ONT) long-read sequencing in conjunction with consensus sequence generation for high-accuracy basecalls. We demonstrate the utility, simplicity, and depth of lineage recording using mouse pancreatic cancer cells (KPCY), ultimately reconstructing single-cell lineage trees with average depths exceeding 29 cell divisions.

**Figure 1.**
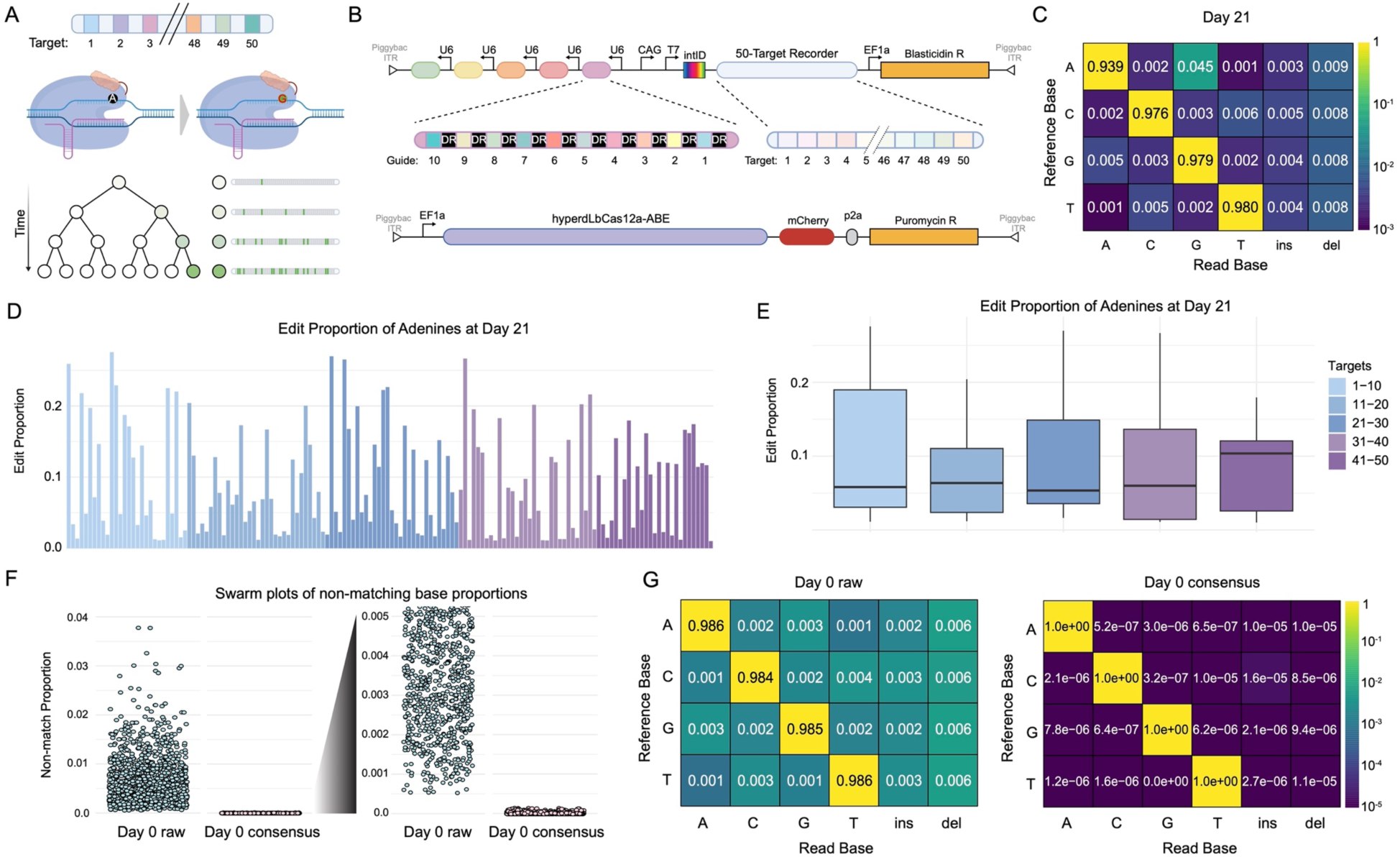
BASELINE records editing events that can be reliably captured with long-read Nanopore sequencing. **A)** Schematic for BASELINE, where adenine to guanine conversions encode lineage information over time. **B)** BASELINE constructs: guides and recorder (top), Cas12a-ABE (bottom). Five guide (gRNA) cassettes each contain a single U6 promoter upstream of 10 multiplexed LbCas12a gRNAs. In the reverse orientation, the 50-target array is directly downstream of the CAG and T7 promoters and the intID (static barcode) sequence. Cas12a-ABE is introduced into cells using either lentiviral transduction or piggyBac electroporation (pictured). **C)** Heatmap conversion matrix showing the proportion of sequenced bases concordant with the reference base at day 21 of editing. Canonical ABE (A-to-G) transitions are shown at the highest conversion proportion. **D)** Full-length recorders revealed 150 edited ‘A’ positions > 1% at day 21 of editing. Different color blocks designate the target sets separated by the five multiplexed guide arrays. **E)** Box plots of editing rates across multiplexed units reveal that all arrays show editing at day 21. Different color blocks designate the target sets separated by the five multiplexed guide arrays. **F)** Box plots that show the total mismatch proportion of raw nanopore reads compared against the mismatch proportion of consensus sequences. On day 0, we expect an edited proportion of zero. **G)** Heatmap conversion matrices show the day 0 raw and consensus sequence nucleotide error rate. Error rates decrease from ∼1 in 1000 to ∼1 in 100000.

## Results

### Building BASELINE

Base editing outcomes are binary, unidirectional changes created without inducing the double-stranded breaks of wild-type CRISPR enzymes^16^. This drastically reduces the number of large deletions observed in previous lineage recording technologies^8,10,27^. To build our initial recorder, we screened computationally generated Cas12a target sequences containing between 3 and 9 editable nucleotides (adenine for ABE; cytosine for CBE) against potential off-target sites in the genome, selecting a library of 990 target sequences (and 10 computationally negative hits). We then tested the editing rates of each target using six Cas12a base editing constructs: three CBEs and three ABEs (**Supplemental File 1, Supplemental Fig. 1A, B**). Across the eight experimental conditions (two controls + six base editors), we recovered an average of 99.175% of our target sequences (**Supplemental Fig. 1C**). We observed the broadest base-editing activity with pSLQ11438, an ABE construct that edited 651 target sequences (**Supplemental Fig. 1D**). Of the CBE base editing plasmids we found that construct 114082 edited the most broadly (252 edited targets). We selected a final target set that maximized editing with ABEs (30 targets), both ABEs and CBEs (19 targets), and CBEs only (1 target) (**Supplemental Fig. 1F**). Of these 50 BASELINE targets, 40 of them are among the most highly editable by ABE (**Supplemental Fig. 1E**), with 272 total adenines sites, of which 147 are within the high-activity Cas12a base editing window^28^ (**Supplemental Fig. 1G**).

To test the feasibility of BASELINE, we designed a minimal recorder with 10 targets and tested the recording with a single multiplexed 10-guide cassette. We stably transfected the 10-target array into 293T cells and transiently transfected in our ABE and our multiplexed guide cassette (**Supplemental Fig. 2A**), where we detected an increased editing proportion across the most highly edited base in all 10 targets. We observed a maximum of 15-fold enrichment of A-to-G editing in the edited condition versus the unedited at the most highly edited sites (**Supplemental Fig. 2B**), establishing the viability of Cas12a-ABE lineage recording in vitro.

Next, we concatenated all 50 of our selected targets into the full-length target array to create large-scale lineage recordings (**Fig. 1A**). Targets and PAMs were placed head to tail without spacer sequences to minimize the recorder length, and multiplexed 10-gRNA cassettes were designed and integrated into a single construct, BASELINE (**Fig. 1B**). We transfected the resulting construct and transduced lentivirus from pSLQ11438 into the KPCY cell line, an established model of pancreatic ductal adenocarcinoma. After 21 days, 4.5% of all recorder adenines had been converted into guanines (A->G), with background error rates between 0.1% and 0.9%. Based on these error rates, we set a threshold (1%) for any nucleotide conversions and found 150 positions where adenines were converted to guanines above the threshold (**Fig. 1D**). When grouped by target, our results showed that all multiplexed guide cassettes were active and producing sgRNA’s compatible with Cas12a base editing, with variable editing rates across the 150 positions (**Fig. 1E and Supplemental Table 1**). To lower our background error rate, we implemented a computational consensus-building approach (**Supplemental Fig. 2C**). Using custom tagging and amplification oligos along with an adapted pipeline for generating consensus sequences^29,30^, we generated consensus sequences with error rates of Q40 (∼1 in 10000) (**Fig. 1F and G**).

### Encoding cellular memories into murine pancreatic cancer cells

We next tested if BASELINE could encode lineage information over time. We cloned the base editing transgene cassette from the lentiviral construct into a piggyBac backbone to compare the difference in the editing accumulation and/or the rate of transgenic silencing and variegation using both systems^31–33^ (**Supplemental Fig. 3A**). We transfected and transduced KPCY BASELINE cells with either the piggyBac-ABE or lentiviral-ABE constructs and quantified the editing proportions after 21 days. Sequencing results showed a higher average editing proportion and more edited sites using the piggyBac-ABE construct (**Supplemental Fig. 3B**). At the least stringent editing threshold of 0.001 (0.1%), we observed 220 sites edited in the piggyBac sample and 183 sites in the lentivirus sample, with average editing proportions at 0.160 and 0.067 respectively (**Supplemental Fig. 3B**). At the most stringent threshold (1%), 157 sites were edited in the piggyBac compared to 129 in the lentivirus sample (average editing of 0.223 and 0.094). This higher activity led us to use the PiggyBac construct in subsequent experiments.

Next, we sought to test whether the editable adenine sites could record lineage independently or if editing at one site would inhibit editing at other adenines within the same Cas12a target^34–37^ (**Supplemental Fig. 4A**). We engineered sequences corresponding to the first 10 targets with their most common single base-edit from our 21-day timepoint dataset (**Supplemental Fig. 4A, Supplemental File 2**). We then generated a stably-integrated KPCY ‘pre-edited’ cell line and exposed these cells to transient base editing. In our 10-target recorder, most alternative sites were still actively edited in our pre-edited sample, at lower rates than in the wild-type recorder (**Supplemental Fig. 4B, C**). Our results recapitulate the findings from other groups^34,38^, showing that a single base pair change does not entirely inhibit Cas12a binding, ensuring independence between neighboring adenines. This data also indicates further editing happens more slowly, reducing the likelihood that a given recorder will become saturated.

We then performed a timecourse experiment where we captured genomic DNA from KPCY BASELINE cells stably transfected with piggyBac-ABE on days post-transfection 0, 7, 12, and 21 (**Fig. 2A**). Across the length of the recorder, we see steady increases in guanine accumulation from day 0 to day 21 (**Fig. 2B**). While the A-to-G changes are most prominent, we also observe the accumulation of T-to-C edits, where the base-editor edits the complement-strand adenines, albeit at a 45X lower rate (**Supplemental Fig. 5A**). Non-canonical editing and deletions increase over time but remain rare (< 1/1,000 bases). At the target level, we also see a uniform accumulation of editing, except for targets 16, 20, and 26 which show a slower accumulation (**Fig. 2C**). The per-target and per-base editing proportion panels follow similar patterns, with the accumulation of edits increasing over time: the difference in day 0 to day 7 is 1.8%, while the difference in day 7 to 14 was 16.7% and day 14 to day 21 was 26.6%, likely due to incomplete antibiotic selection at earlier time points.

**Figure 2.**
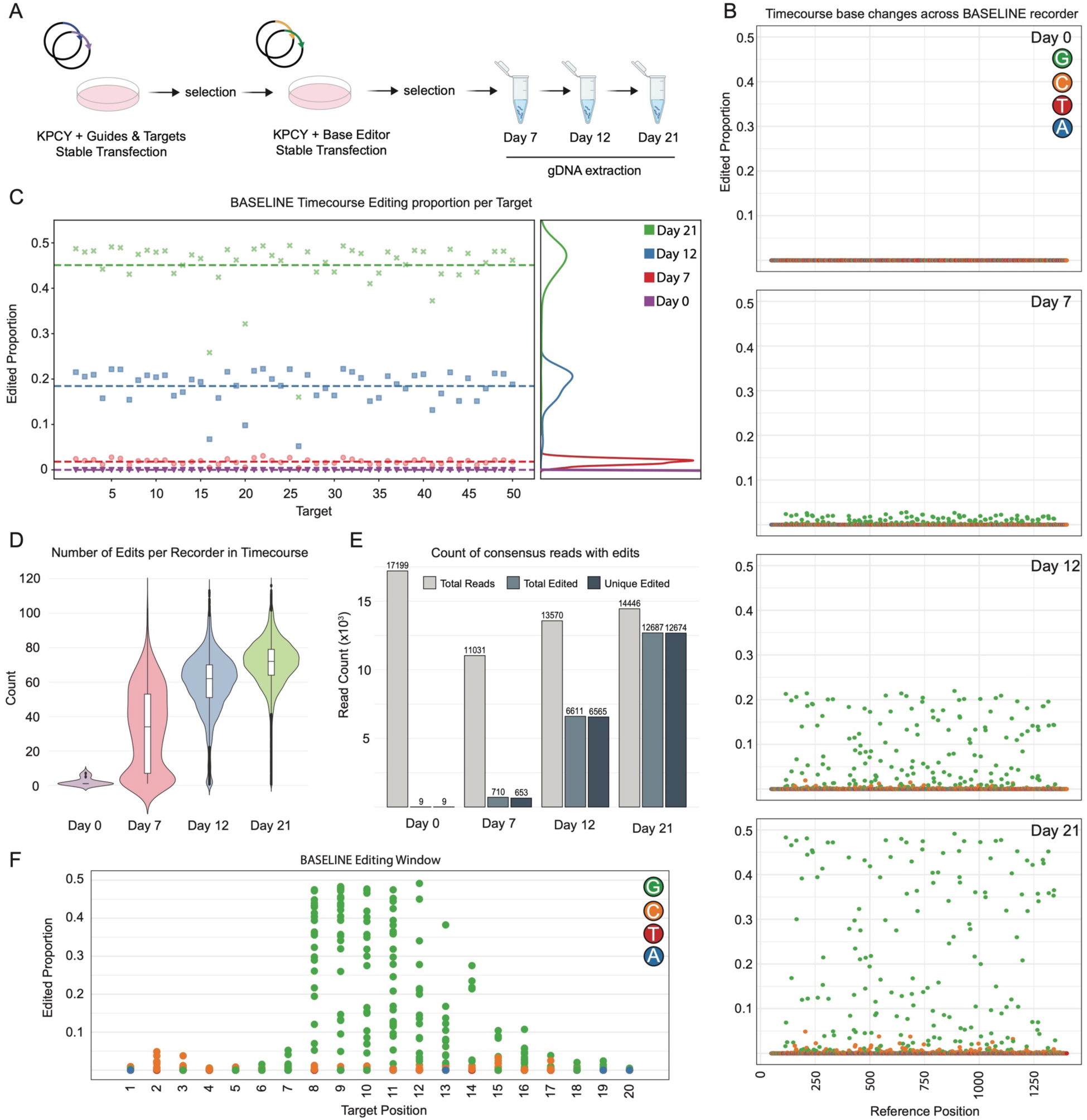
Timecourse experiment showing temporal and canonical editing accumulation. **A)** Workflow schematic for the time course experiment. **B)** Dot plots of each timepoint (day 0, 7, 12, and 21) showing the per-base changes across the entire recorder length. Green = G, Orange = C, Red = T, and Blue = A. **C)** Dot plot and paired density plot show the per-target editing proportion increases through time. Purple = Day 0, Red = Day 7, Blue = Day 12, and Green = Day 21. **D)** Violin and paired box plots show the edit count of edited recorders (colors as in panel C). **E)** Bar plots showing the total number of consensus reads (light-grey), the total number of edited reads (slate), the total number of uniquely edited reads (navy) for each time point in the timecourse. Consensus sequences for each time point were filtered by intID and editing pattern to remove duplicates. **F)** Dot plot that shows the accumulation of guanine edits along the length of all BASELINE targets at day 21, showing the active editing window.

Lastly, we investigated editing pattern variability across integrations. Most recorders were uniquely edited (**Fig. 2E**), and unique editing patterns were also seen within each target site (**Supplemental Fig. 5B, Supplemental File 4**). In target 8, with 32 (2^5^) potential outcomes, nine appear in greater than 1% of the reads. We can expand this to all 50 targets and use the product of editing patterns to find that our total information recording capacity exceeds 1.90*10^35^ (**Supplemental Table 2**). While this capacity is likely inflated due to the variable editing rate at different target adenines, it still exceeds the necessary capacity for mammalian phylogenetic tree reconstruction (∼3.7*10^12^)^39–41^. In this time course experiment, BASELINE accumulates edits constantly over the 21 days, edits are spread across all targets of the recorder, and the recorders accumulate edits in a pattern that makes them unique and distinguishable from other edited recorders.

### Simulations of BASELINE reconstruct highly accurate phylogenetic trees

Given the editing rates detailed above, we then set out to measure the reconstruction accuracy of our BASELINE lineage approach. We devised a computational framework to simulate lineage trees using both our observed editing rate (back-calculated to a per-generation rate, see methods) and a range of potential editing rates (**Fig. 3A**). We included a dropout rate to simulate the challenges of recovering the recorder with single-cell sequencing technologies. We were also interested in how sparse sampling of a large lineage tree would affect reconstruction accuracy, for instance, subsampling 5,000 cells from a whole organismal lineage tracing experiment. To that end, we generated lineage trees ranging in size from 2^4^ (16) to 2^20^ (∼1 million cells), randomly sampling 2^12^ (4,096) cells from each tree. Lastly, we varied the number of integrated recorders from 1 to 30, representing the range of integrations observed in our KPCY and 293T models.

**Figure 3.**
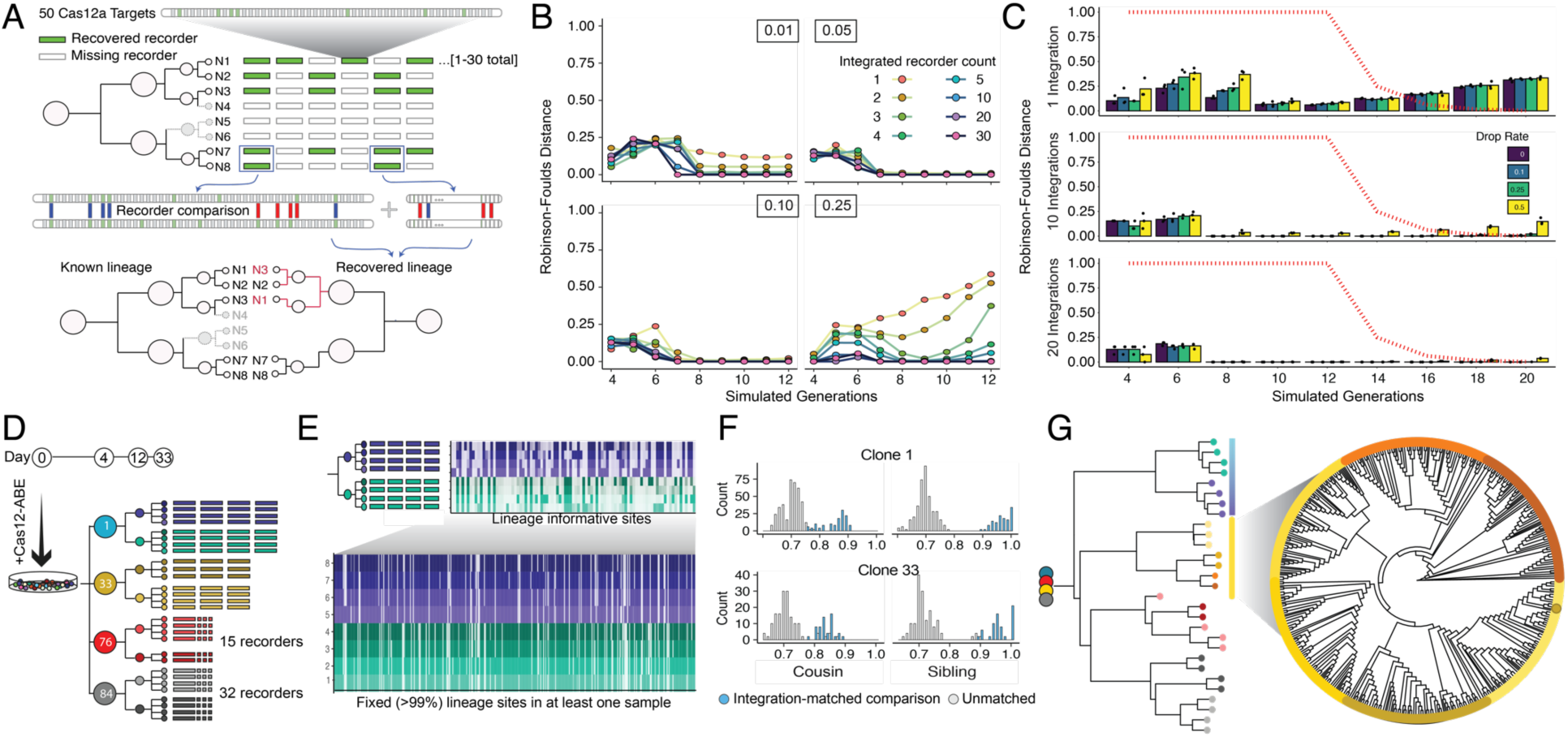
In silico and in vitro lineage simulations recover accurate trees. **A)** In silico lineage reconstruction using 50-target lineage recorders. To account for the challenge of recorder recovery, we then randomly dropped from 0-50% of each cell’s recorders. Trees were then reconstructed, and accuracy was computed using Robinson Foulds’ distance to the known simulation tree. **B)** Simulation accuracy from 4 to 12 generations for 10% recorder dropout rate over a range of integrated recorder counts and editing rates (inset box). **C)** Simulated lineage reconstruction distance using editing rates established in the previous timecourse experiment from 4 to 20 generations with dropout rates from 0 to 50%. Red lines indicate the proportion of total cells sampled at each time point. **D)** Synthetic lineage created using cell culture; Single BASELINE KPCY cells were isolated from a polyclonal population, and individual cells were isolated from these cells twice over, creating granddaughter populations with known lineage relationships. **E)** In clonal population 1, 367 A>G sites were fixed (>99% edited, bottom) in at least one granddaughter population (of 1089), where 75 were fixed in one population and absent (<1%, top) in another. **F)** Recorder correlation increases from unrelated recorders (r=.74) to integration-matched recorders in cousins (0.87) and siblings (0.94), capturing lineage information. **G)** Lineage reconstruction using hierarchical clustering of granddaughter wells, blinded to known lineage relationships, and in silico single-cell generation of single-cells accurately reconstructed lineage relationships between 600 simulated granddaughter cells.

Each simulation was run in triplicate, and trees were reconstructed using IQ-Tree^42^. We then compared these reconstructed trees to the simulated ground-truth trees using the normalized Robinson-Foulds (RF) distance^43^. Setting the per-generation editing rate to specific values revealed higher accuracies at moderate editing rates, with improved performance at higher integration counts (**Fig. 3B, Supplemental Fig. 6A**). We achieved accurate (∼100%) reconstructions at 12 generations, with the lowest reconstruction accuracies observed with high per-generation editing rates, driving early saturation of the recorders (**Supplemental Fig. 6A**).

We then simulated trees using the editing rates established in our KPCY time course (**Fig. 3C, Supplemental Fig. 6B**), where we again obtained high-accuracy reconstructions across most parameters. Despite even sparse sampling of individual cells (0.3% of cells recovered), we recovered 95% accurate lineage trees with three recorder integrations. We achieved ∼100% accuracy with ten or more integrations (**Fig. 3C, Supplemental Fig. 6B**). The most inaccurate reconstructions occurred when using <5 recorders, a 50% recorder dropout rate, and at early cell divisions, causing accuracy to drop to ∼50% in the three-recorder, six generation simulations (**Supplemental Fig. 6B**). In these simulations a single recorder often outperformed two to five recorders, likely single recorders are directly comparable, as opposed to the low cell-to-cell comparison rate in two-or three-recorder simulations.

We generated an on-plastic lineage tree using our in vitro KPYC system to further validate BASELINE’s ability to record lineage relationships. We electroporated KPCY cell populations with our construct and isolated individual clones, which we characterized for the number of integrations using our nanopore pipeline. High-confidence integration counts ranged from 3 to 32, from which we selected four populations to expand into a synthetic lineage tree (**Fig. 3D**, **Supplemental Table 3**). We initially focused on clone 1, containing four integrated recorders, which was successively split into eight granddaughter populations. Of the 1,085 sites across the four integrated recorders, 364 were fixed (> 99% edited) in at least one of the granddaughter populations (**Fig. 3E, bottom**). Of these, 46 sites were fixed in at least one population and absent (< 1%) in another, indicating a lineage-informative site (**Fig. 3E, right**). 24 of these sites statistically differed between the parental groups (Wilcox test < 0.05).

To ensure that BASELINE recorded lineage-specific information at all levels of the resulting tree, we compared the correlation of base editing outcomes across cousin populations (sharing a grandparent) and within sibling populations (**Fig. 3F**). In clonal population 1, sibling cell recorders were >90% correlated, which decays to 87% in cousin cells and to 74% in unrelated cells, representing lineage specificity over the background of shared editing patterns (**Fig. 3F**). A similar relationship is seen in clone 33 data.

We used these aggregated editing proportions in each granddaughter population to test tree reconstruction. We used average editing rates at each integration to create a population-level recorder for each granddaughter well, which was reassembled into a population-level tree and faithfully reconstructed clonal relationships (**Fig. 3G**). Although our bulk DNA sequencing data does not allow us to assign recorders to individual cells, we were curious if aggregating recorders into synthetic cells would still enable accurate tree reconstruction. We simulated drawing 600 cells from the set of integrations in each of the clone 33 daughters (3 integrations per cell). Across 100 simulated trees, 99.73% of simulated cells were assembled closest to a sibling cell on the resulting tree, with very sparse misassignment of cells to cousin populations, an example of which is shown in **Fig. 3G**. Our simulations demonstrate BASELINE’s high recording capacity and accuracy (>99%) with real-world base-editing rates.

### Accurate single-cell reconstruction of KPCY clonal populations

To understand BASELINE dynamics in vitro and test lineage recorder expression and recovery, we subjected KPCY BASELINE cells to single-cell RNA sequencing (scRNAseq) on the 10X Genomics platform. We piloted three approaches to maximize the recovery of BASELINE recorders: (1) capture-sequence driven recovery^44^, (2) hybridization-capture, and (3) in vitro transcription (**Supplemental Fig. 7**). The capture-sequence (CS1) approach led to unintended products and required excessive amplification; hybridization capture improved the recovery of the intended band but still suffered from unintended products and required over-amplification (**Supplemental Fig. 8**). In vitro transcription (IVT) created clean product bands, required fewer than 10 cycles of PCR amplification, and allowed us to simultaneously isolate BASELINE recorders captured from the oligo(dT) and the 10X capture sequence pools (**Supplemental Fig. 8**).

With an established recorder recovery and isolation protocol, we selected two clonal populations for lineage reconstruction: clone 76, which ultimately contained 19 integrations, and clone 84, with 34 integrations (**Supplemental Fig. 9A**). Both clones were transfected with piggyBac-ABE on day 0 before being sorted as single cells into individual wells of a 96-well plate on day 4. We expanded and passaged the clones to confluency in a single well of a 6-well plate (day 19), which we sequenced using the 10X Chromium (**Fig. 4A**). After recorder recovery and isolation, we sequenced BASELINE recorders on a PromethION flowcell and single-cell transcriptomes on an Illumina NextSeq. The transcriptional profiles of clones 76 and 84 showed divergence from each other and an additionally sequenced parental KPCY population (**Fig. 4B**), highlighting transcriptional states observed in our previous work^10^.

**Figure 4.**
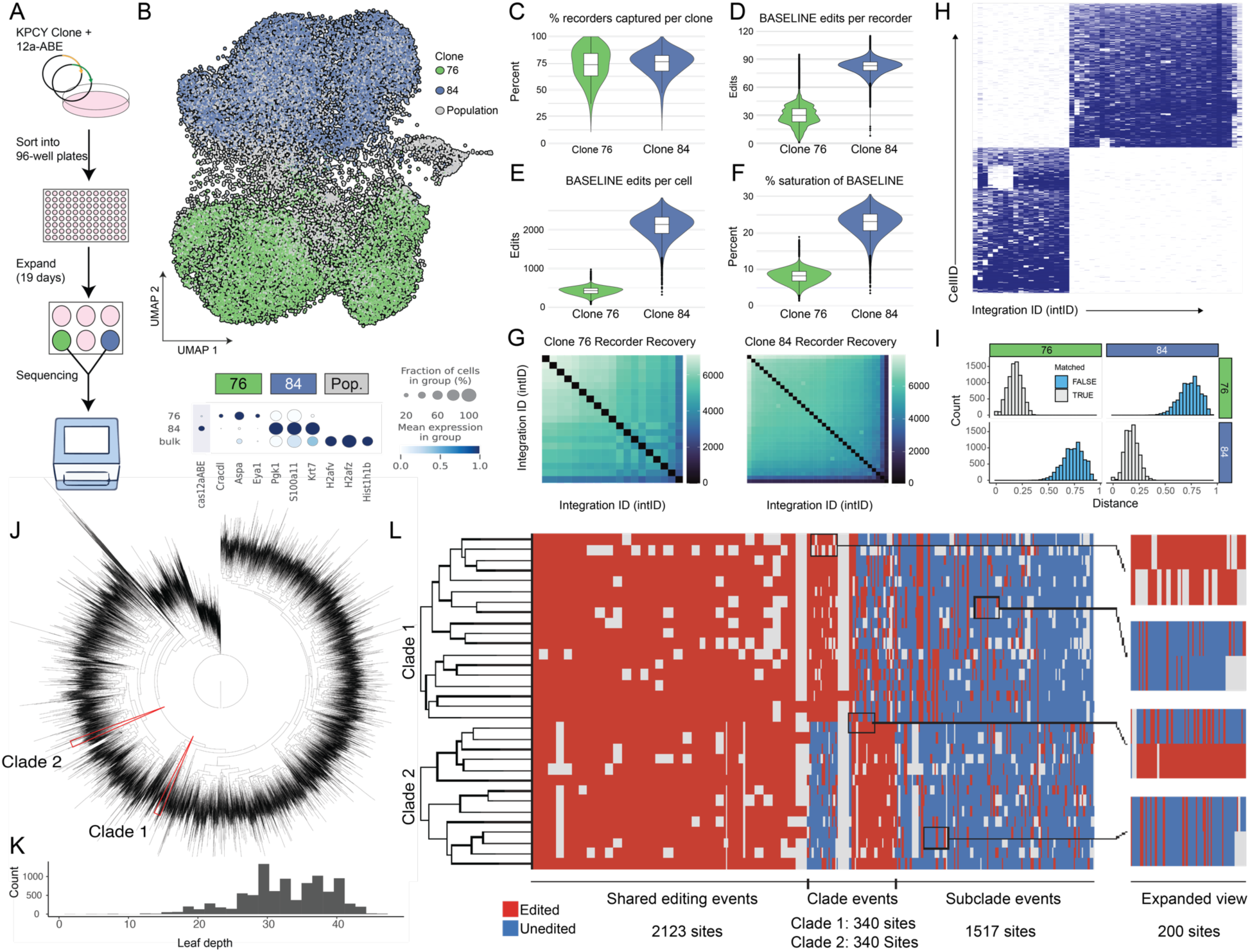
Single-cell sequencing of BASELINE yields efficient recorder capture, editing, and reconstruction of phylogenetic trees. **A)** Experimental workflow for clonal expansion and droplet-based scRNAseq of BASELINE clones 76 and 84 from isolated single-cell populations. **B)** Single-cell sequencing of 24,807 KPCY cells from the control population (gray), clone 76 (green), and clone 84 (blue). Expression of Cas12a-ABE and marker genes (bottom) **C)** Capture rate per cell per clone, **D)** the number of edits per recorder per clone, **E)** the number of edits per cell, and **F)** the percent saturation of the cells within each clone. **G)** Heatmaps of recorder integration recovery within clones 76 and 84. **H)** Hierarchical clustering of integration IDs (intIDs) and 18,441cellIDs from the combined clone 76 and 84 data. Clone 76 contains 19 total intIDs, and clone 84 contains 34 intIDs. **I)** Similarity comparison of permuted recorders from clone 76 and 84. **J)** Phylogenetic tree reconstruction for clone 84. Two clades are outlined in red arrows, which are expanded in panel L. **K)** Distribution of leaf depths in clone 84 tree reconstruction. **L)**. Tree substructure from panel J. Clade 1 contains 18 cells while clade 2 each contain 14 cells. Of 9248 adenines, both clades share 2,123 edited sites. Clade 1 contains 340 fully edited sites not edited in clade 2; clade 2 contains 340 unique sites. 1,517 sites vary within the clonal population, and the remaining (>4900) remain unedited or uncaptured in both.

We then reconstructed individual BASELINE transcripts using a custom pipeline based on the scNanoGPS toolset^45^. Analysis of the 10X cell barcode (cellID) recovery showed that 18,776 (99.64%) of cellIDs identified by CellRanger and processed with Seurat^46^ were also recovered by scNanoGPS (**Supplemental Fig. 9C**). These overlapping cellIDs were assigned to a clone if ≥90% of their captured intIDs belonged to that clone (**Supplemental Fig. 10**), leaving us with 18,441 cells (9,260 for clone 76 and 9,181 for clone 84). Our median recorder capture per cell was 73.68% and 76.47%, recovering ∼14/19 and ∼26/34 recorders per cell for clones 76 and 84 respectively (**Fig. 4C**). The median base-edit count per recorder was 30 for clone 76 and 83 for clone 84 (**Fig. 4D**), with a median of 427 and 2,140 edits per cell (**Fig. 4E)**, representing a total adenine saturation percentage of 8.26% and 23.14% (**Fig. 4F**). In clone 84, greater than 75% of cells contained over 1,900 edits, a 50-fold increase over the state-of-the-art^47^. We filtered out any recorder with an ambiguous basecall to find that over 97.8% of the recorders captured in clone 84 had a unique editing pattern (**Supplemental Fig. 12)**. We then calculated the editing percentage for each of the 272 adenines (**Supplemental Fig. 13A, B)**. Clone 84 had 9 positions with a saturation percentage over 99% at 19 days and 36 positions over 90%, leaving significant recording capacity for future use. Clone 76 showed considerably lower editing percentages across all adenine positions and had no positions saturated in all recorders.

With this large set of recorders, we were curious how capture efficiency varied across integration sites. Recovery was remarkably uniform, with lower rates in two integrations per clone and little correlation structure between integrations (**Fig. 4G**). Hierarchical clustering of the filtered cells and the known intIDs showed clear separation and capture purity for clone 76 and 84 (**Fig. 4H, Supplemental Fig. 9B, 11**). A subpopulation of clone 76 cells exhibited a common pattern of failed recorder captures (1,237 cells missing recorders 1, 4, 5, 6, 7, and 8), suggesting a higher degree of relatedness. A genomic event like transposon eviction or chromosome loss followed by clonal expansion could explain this shared lineage phenomenon. We also compared editing distances between randomly sampled recorder pairs from clones 76 and 84 and showed a strong correlation between within-clone recorders and substantial divergence between clones 76 and 84 (**Fig. 4I**).

Using IQ-TREE, we reconstructed phylogenetic trees from clone 76 and clone 84 (**Fig. 4J, Supplemental Fig. 14**). The reconstruction of clone 76 resulted in an asymmetric tree caused by the low levels of editing throughout the cell population. Differential gene expression analysis of our KPCY populations showed low Cas12a-ABE expression in clone 76, contributing to the low resolution (**Fig. 4B, Supplemental Table 4**). Despite low Cas12a-ABE expression, we mapped the 1,237 cellIDs from our hierarchical clustering lacking recorders 1, 4, 5, 6, 7, and 8 onto the phylogenetic tree (**Supplemental Fig. 15**). We observed that >98% of these cellIDs map to a single tree clade, validating lineage reconstruction. This co-clustering was preserved when we excluded these missing recorders from all cells (**Supplemental Fig. 16**).

We achieved a more uniform recovery of clone 84 recorders (**Fig. 4H)**. These constructs had highly unique outcomes, with an average depth of 29 cell divisions and maximum depths of 40 (**Fig. 4K**), and with 91.7% of tree leaves with ≥ 20 cell divisions. We sampled two clades to profile the support at each branch point and determined the support for their unity and divergence (**Fig. 4L**). 2,123 sites were jointly edited in both clades, 340 uniquely supported the substructure in each clade, and 1,517 sites were partially edited in a subclade, representing local lineage structure. Clone 84 then retains 4,931 sites for future lineage recording across both clades.

## Discussion

This study introduces BASELINE, a lineage tracing tool capable of reconstructing accurate phylogenetic trees over biologically relevant timescales. Our Cas12a-based system has several advantages over Cas9 systems, most importantly, the ability to multiplex large numbers of guide sequences and targets into compact payloads^48,49,50,51^. Additionally, our base-editing system avoids large intra-recorder deletions, resulting in the loss of lineage information^52^. We demonstrate reliable recovery of the 1.5 kbp recording array using IVT, and we leverage ONT long-read sequencing to generate accurate consensus sequences alongside single cell transcriptomes.

BASELINE recorders are well-expressed and captured in scRNAseq; we recovered at least one recorder in ∼100% of cells and the median cell recovered ∼75% of all possible recorders. Our single-cell lineage trees show the advantages of this large recording space; in the clone 84 lineage tree, we achieved an average tree depth of 29 cell divisions, with ∼4,900 remaining base edits for future recording. In clone 76, we validate tree reconstruction with a shared chromosomal event, correctly assigning >98% of cells. Together, the features of BASELINE form a system that can exceed the current state of the art while simplifying lineage tracing engineering.

The ultimate aim of lineage technologies is to capture lineage maps of every cell division in conjunction with single-cell state. While our results indicate that BASELINE represents an advance in the state-of-the-art, there are limitations. As designed, BASELINE needs at least two integrations per cell to reconstruct lineage trees with 92% accuracy after 12 generations. With our current antibiotic selection-based constructs, single-cell isolation and screening must be used to select high-integration founder cells. Our modifications that allow for a median recorder recovery of nearly 75% shows substantial progress in improving transcript capture and expression; however, 100% recovery remains the goal. As has been shown by our simulations, the editing rate affects phylogenetic reconstruction and, despite capturing over 9,000 cells each from two clonal populations, our variable editing rate was reflected in clone 76’s more asymmetric reconstruction. While our system relies on the assumption that Cas12a remains constitutively active, expression-silencing mechanics could limit recording capacity and significantly impact our ability to reconstruct lineage trees.

Despite these limitations, BASELINE is an efficient, simple, and viable option for any group interested in lineage tracing experiments. We use Cas12a-ABE, building upon previous efforts^53^, leaving Cas9 open for gene perturbation or as an additional recording channel. Additionally, base-editing approaches are compatible with single-cell transcriptional and hybridization-based spatial sequencing^54^. With high reconstruction accuracies across a range of recorder integrations, advancements to long-read sequencing technologies, and improved transcript capture for scRNAseq methods, we foresee continued progress toward biological insights with BASELINE.

## Data Availability

Raw sequencing data has been uploaded to the Sequence Read Archive with the associated BioProject ID PRJNA1224774. Single-cell sequencing data has been uploaded to the Gene Expression Omnibus with the associated BioProject ID GSE289815.

## Methods

***(all product and kit-based reactions were performed as per the manufacturer’s instructions unless otherwise stated)

### Cell lines culturing and maintenance

All experiments were carried out in HEK293T (ATCC CRL-3216) or KPCY (Kerafast PDAC 6419c5). HEK293T cells were cultured in DMEM (Corning™ 45000-316) with 10% FBS and passaged at a ratio of 1:5 every two days. KPCY cells were cultured in DMEM (Corning™ 45000-304) with 10% FBS and passaged at a ratio of 1:10 every three days with a media change on day two.

### Plasmid Construction

All plasmids were built with NEBuilder® HiFi DNA Assembly Master Mix (NEB E2621L) according to the manufacturer’s protocols. All plasmids were sequenced and confirmed using Circuit-Seq^55^. The BASELINE plasmid was built into the backbone of the piggyBac donor vector. The original donor vector was cloned to replace the puromycin resistance with blasticidin resistance (PB-blast). PB-blast was digested with SpeI and BamHI and gel extracted to retain the long fragment. The CAG promoter was digested from a separate donor vector and the 50-target recorder was produced by IDT as a gBlock and both were ligated into the cut PB-blast (PB-50T). The five separate ten-guide cassettes were produced with oligos containing protospacers and direct repeats were hybridized and extended to create dsDNA fragments that overlapped with neighboring dsDNA protospacer fragments (**Supplemental File 1**). Each 10-guide DNA fragment was ligated into pLJM1-EGFP-U6. Each guide cassette was amplified from their respective construct with appropriate oligos (**Supplemental File 2**). PB-50T was cut with StuI for golden gate assembly with five amplified guide cassettes (PB-50G-50T). PB-50G-50T was then cut with SwaI for insertion of the T7 promoter and a drop-in site for the static integration identifier (PB-T7-50G-50T). Degenerate intIDs were inserted into PB-T7-50G-50T after cutting with KpnI (BASELINE).

The pLJM1-EGFP-TWIST backbone was built into the pLJM1-EFGP construct. The U6 promoter from pRDA-052 was excised and ligated into the backbone. The standard drop-in site was removed and replaced with either the AsCas12a DR or the LbCas12a DR sequence along with a PaqCI drop-in site. Nuclease plasmids were purchased from Addgene (**Supplemental File 1**).

PiggyBac Cas12a was built into the backbone of the piggyBac empty vector. The empty vector was cut with SpeI and NheI and the CMV promoter was discarded. The remaining fragment was ligated together and cultured before being digested with EcoRI and SalI to keep the 4.9 kbp fragment. Addgene pSLQ11438 was digested with EcoRI and SalI and the 7.5 kbp fragment containing the expression cassette was ligated into the piggyBac backbone.

The re-editing experiment required the production of two constructs. Each construct was built with a 10-target array made from IDT as a gBlock and the backbone of the PB-T7-50G-50T construct. PB-T7-50G-50T was digested with StuI and PmlI to remove the CAG promoter and 50-target recorder. The gBlocks were separately ligated into the backbone (PB-10T-preedit and PB-10T-unedit).

### TWIST oligo pool

10,000 target sequences (20nt in length with TTTC PAM motifs) were generated with FlashFry^56^. These sequences were analyzed using the CRISPR/Cas target selection tool: http://www.rgenome.net/cas-designer/. The top 990 predicted target sequences along with 10 non-targeting control sequences were generated as an oligo pool by TWIST Biosciences. Oligo design can be found in **Supplemental File 3**. Oligos were amplified using ETC and ETD and were digested with PaqCI for drop-in to PaqCI digested pLJM1-EGFP modified.

### Lentivirus

Production of lentivirus was performed by co-transfecting pMD2.G, psPAX2, and the transfer plasmid into a 10 cm dish of 293T cells. Media was changed at 24h, 48h, 72h, and 96h. Media was kept from timepoints, 48 h, 72 h, and 96 h and filtered with a 0.45 µm syringe filter. Filtered media was concentrated with the Sorvall WX80+ Ultracentrifuge. Concentrated lentivirus was resuspended in DMEM complete media and stored at -80°C.

### Chemical Transfection

Plasmids transfections were performed with PEImax (1 mg/mL) at a 3:1 ratio of PEI:DNA. For a 24-well transfection 500ng of plasmid was mixed with 25 µl of Opti-MEM SFM. 1.5 µl of PEI was added dropwise to the solution and mixed by gently flicking the tube. After a 10-minute incubation at room temperature, the final mixture is added dropwise to the 24-well for a total of 300 µl media and transfection solution. Media was changed after 24 h to 500 µl complete DMEM. 24-well was split into 12-well after 48 h and selection was added at 72 h.

### Electroporation

Plasmid electroporations were performed with the Neon™ Transfection System according to the manufacturer’s instructions. KPCY cells at a concentration of 5.6*10^6^ were mixed with a total of 5 µg plasmid and electroporated with conditions: 1,150 V, 30 ms, 2 pulses. Electroporated cells were plated into one well of a 6-well plate. At 24 h media was changed and included selection.

### PCR Amplification and Clean-Up

All amplification reactions were performed with Q5 Hot Start 2X Master Mix (NEB M0494L) using SYBR^TM^ Green I Nucleic Acid Gel Stain (Thermo Fisher Scientific S7563) and the CFX Connect Real-Time Detection System (Bio-Rad). Amplifications were manually stopped before the plateau phase of the qPCR curve, unless otherwise noted. After amplification, PCR reactions were cleaned using the DNA Clean & Concentrator-5 Kit (Zymo D4014) (for cloning products) or magnetic beads (Omega Biotek Mag-Bind® TotalPure NGS) (for material to be sequenced).

### Next Generation Sequencing

DNA from cell lines was extracted using the DNeasy Blood and Tissue Kit (cat. 69504). 300 ng of genomic DNA was tagged in 2 cycles of PCR with oligo ETQ and cleaned up with magnetic cleanup beads. The elution of tagged amplicons was amplified using ETR and ERU for 20 PCR cycles before being cleaned up again with magnetic beads. Truseq i7 and Nextera i5 combinatorial indexes were added with 4 additional cycles before a final bead cleanup. Libraries were pooled and sequenced on the Illumina MiniSeq device using the mid output 300-cycle kit.

### Oxford Nanopore Sequencing

Separate amplicon samples were barcoded with the Native Barcoding Kit 24 V14 (SQK-NBD114.24) before being pooled and sequenced using a PromethION Flow Cell R10.4.1 (FLO-PRO114M). Flow cells were washed and re-loaded with more libraries every 24 hours until the flow cell was not viable or until no more libraries remained. For the re-editing experiment, oligos EFV and EFT were used to amplify the 10-target fragments before being barcoded as above. The resulting barcoded sample was loaded onto a Flongle Flow Cell R10.4.1 (FLO-FLG114) and ran until no longer viable. All sequencing output POD5 files were basecalled with the Dorado 0.7.0 basecaller and output as FASTQ files.

### Single Cell Dilution Sorting

We acquired images of our 96-well plate at 20X magnification in the RFP channel. These images were analyzed by Fiji to determine the approximate cell count. The chosen wells were trypsinized with 30 µl of 0.25% trypsin (Corning^TM^ MT25053CI). Wells were washed with media and into a total final volume of 1mL media in a microcentrifuge tube. Approximately 0.5 cells per 100 µl were transferred into a reservoir with 25 mL of complete media. Media in the reservoir was gently mixed with a serological pipette 3 times. After mixing, 100 µL of cell-containing-media was dispensed into new 96-well plates. 96-well plates were not disturbed for 4 days until 50 µl of fresh media was added to the wells. On day 7 the wells were scanned for growth and wells with observable cells were given new media.

### Tagging PCR for genomic amplicon consensus (pipeline-umi-nf)

An abbreviated protocol is described below, for the full protocol, please refer to Amstler et al. Extracted DNA was tagged with 4 cycles of PCR using tagged oligos (FVO and FVP). The tagged PCR product was purified with 0.6X magnetic beads. Elution from tagging PCR was amplified for 5 cycles (short PCR) with amplification oligos (FVQ and FVR). Short PCR reactions were purified again with 0.6X magnetic beads. Elution from short PCR was amplified for 20 cycles (long PCR) with amplification oligos (FVQ and FVR). Long PCR reactions were purified with 0.45X magnetic beads and amplicon quality and length were verified using DNA electrophoresis.

### Single-cell RNA sequencing

Clones 76 and 84 were stably transfected on day 0. Antibiotic selection was added on day 2 and continued throughout day 10 of the experiment. On day 4, clonal populations were dilution sorted to keep 1 cell-per-well in a 96-well plate. 10 colonies per plate were expanded until a confluent 6-well plate was obtained (day 19). Colonies with the most uniform RFP expression progressed to the scRNAseq step (1 population for each clone). Cells were prepared as indicated in the 10X GEM-X single-cell preparation protocol. The clone 76 and clone 84 populations were mixed in a 1:1 ratio and diluted to 1,100 cells/ul and a targeted cell recovery of 20,000. The protocol was followed as indicated by the manufacturer. Transcriptome libraries were produced and sequenced on the NextSeq 2000 before being analyzed with Seurat (v5.1.0).

### BASELINE In-vitro Transcription and Library Preparation

The protocol for the Chromium GEM-X Single Cell 3’ v4 Gene Expression with Feature Barcoding (10X Genomics) was followed through step 2. Of the 40 µl cDNA amplification output, 5 µl was sequestered for RNA synthesis with the HiScribe® T7 Quick High Yield RNA Synthesis Kit (NEB E2050). The IVT reaction was incubated at 37°C for 4 hours before being treated with DNaseI at 37°C for 30 minutes. The resulting RNA was purified using the Monarch® Spin RNA Cleanup Kit (NEB T2040L). RNA concentration was quantified using the Implen NanoPhotometer® N50. Following the manufacturer’s instructions, the first-strand synthesis was initiated with 100ng of RNA and the FDZ oligo with the ProtoScript® First Strand cDNA Synthesis Kit (NEB E6300). The oligo(dT) captured transcripts were amplified 1/10th of the cDNA reaction with primers FUI and FDZ. The recorder transcripts captured with the CS1 oligo were amplified using the FUI and EWE oligos. Amplifications were stopped at 5000 RFU (≤6 cycles) and were purified with 0.45X magnetic beads. The eluted product was checked for fragment size and purity with DNA gel electrophoresis. If quantity or purity was insufficient, additional amplification and/or magnetic bead purification were performed.

### Synthetic Lineage Tree

We electroporated KPCY BASELINE monoclonal populations with piggyBac-transposase and PB-Cas12a-ABE (day 0). On day 4, we performed single-cell dilution sorting with 100 µl selection media (2 µg/mL puromycin and 10 µg/mL blasticidin) on the electroporated clones and dispensed them into 3 x 96-well plates. On day 7 we added 50 µl complete media into all wells. On day 12, some wells had >1000 cells and were RFP positive. We used Fiji to count the cells in the RFP-positive wells. We picked two RFP positive wells per clone and dilution sorted again into 2 x 96-well plates. The dilution sorting step was repeated on day 19 for 2 more wells. The growing wells were expanded into 48-well plates and we performed gDNA extraction with DNeasy Blood and Tissue Kit at >95% confluency (day 33).

### Time course Experiment

BASELINE constructs with intIDs were electroporated into KPCY cells with piggyBac-transposase on day -7. Cells were propagated and selected with 20 µg/mL of blasticidin for 7 days post-electroporation. *Electroporation:* On day 0, BASELINE cells were electroporated with PB-Cas12a-ABE and piggyBac-transposase. *Transduction*: On day -1, BASELINE cells were passaged into 8 wells of a 24-well plate at 80K cells per well. On day 0, approximately 20-hours post-passaging the cells were transduced with lentivirus from pSLQ11438. *Both conditions*: An aliquot of day 0 KPCY BASELINE cells were washed and gDNA extracted. KPCY BASELINE cells were propagated and selected with 20ug/mL blasticidin and 2ug/mL puromycin until day 7. On day 7, selection was removed for 3 days before continuing again with 20 µg/mL and 2 µg/mL of blasticidin and puromycin respectively. One million cell aliquots were removed on days 7, 12, and 21 and their DNA was extracted using the DNeasy Blood and Tissue Kit.

### Re-Editing Experiment

KPCY cells were electroporated with either of the two target-containing constructs (PB-10T-preedit or PB-10T-unedit), with PB-Cas12a-ABE, and with piggyBac-transposase. Cells were selected with 20 µg/mL blasticidin and 2 µg/mL puromycin until day 7 at which point selection levels were reduced to 10 µg/mL and 1 µg/mL for blasticidin and puromycin, respectively. One million cell aliquots were removed on days 7 and 14, and their DNA was extracted using the DNeasy Blood and Tissue Kit.

### Hybridization Capture and Library Preparation

The protocol for the Chromium GEM-X Single Cell 3’ v4 Gene Expression with Feature Barcoding (10X Genomics) was followed through step 2. Of the 40 µl cDNA amplification output, 5 µl was sequestered for amplification with oligos EWI-biotin and FDZ. The PCR product was purified with 0.45X magnetic beads. Eluted DNA was carried forth using the Tube Protocol from the High-throughput hybridization capture enrichment of long genomic fragments (IDT xGen^TM^ Hybridization and Wash Kit v2). After heated and RT wash steps the beads are moved to the amplification step. Using oligos EWI (not biotinylated) and FDZ amplification is run for 10-15 cycles. Amplification reagents are provided in the hybridization capture kit. The resulting product was separated from the streptavidin beads using a magnet. The PCR products were analyzed for purity and length using DNA gel electrophoresis (**Supplemental Fig. 8D and E**).

### Analysis for umi-pipeline-nf

Consensus sequence generation utilized the tool umi-pipeline-nf^29^. We followed the tool manual that can be found on github (https://github.com/genepi/umi-pipeline-nf). A version of the config file that we used can be found on github (https://github.com/ebwinter95/BASELINE). The parameters that we changed for BASELINE were: min_read_length, min_reads_per_cluster, min_consensus_quality, and medaka_model.

### Analysis for scNanoGPS

To generate consensus sequences of the ONT sequenced IVT product, we used the tool scNanoGPS^45^ (https://github.com/gaolabtools/scNanoGPS). Before running scNanoGPS we preprocessed the reads. First, we aligned the reads to the reference sequence (minimap2 2.24-r1122). Then we filtered the aligned reads by their AS score and their sequence read length. After, we run the Scanner script. Changes were made to the default settings for the 5’ adapter (GCTAGACATTGTGCCGCATC) and the scanning region length (200). We then run Assigner script with a forced cell number at 23,000. We then ran the extract_reads_for_scNano.py script to filter out cell recorders with fewer than 20 associated UMIs (manually inspect the CB_counting.tsv.gz output from assigner.py to obtain the number of CBs with more than 20 UMIs). This code also limits the number of CB-UMI pairs to 20 sequences per pair (this keeps the consensus-building step from freezing due to generating a consensus with an overwhelming number of sequences). Lastly, this code requires a CB-UMI pair to appear at least 5 times for the reads to be included in downstream analysis. We used the extracted read names to filter the aligned and AS-filtered bam file. Finally, with our preprocessed reads, we re-ran the Scanner tool and re-ran the Assigner tool (we used the forced cell number found from inspecting CB_counting.tsv.gz from the first run). Then, we ran the Curator tool with the keep-meta option set to ‘1’ and with edits to the curator_io.py file. We then ran the convert_sam_to_bam.py script before running test_code_from_chatgpt.py, process_fastas.py, and binary_df_maker-Copy1.py (for clone 84) and binary_df_maker.py (for clone 76). The non-scNanoGPS scripts, edited curator_io.py file, and a protocol for running this single-cell analysis can also be found at https://github.com/ebwinter95/BASELINE/tree/main/running_scNanoGPS.

### Cell Lineage Simulation

All cell simulations were performed with Counterfeiter, a framework for simulating lineage trees (https://github.com/mckennalab/counterfeiter). Counterfeiter was either given (1) the established base-editing results from the timecourse experiment, in which we approximated the per-cell, per-generation editing rate, or (2) set editing rates. For the existing in vitro editing rates, we established a per-generation rate. Given the unidirectional nature of base-editing and the interdependence of editing rates over the tree, we can approximate a per-cell editing rate by calculating the chance of 1 minus not observing any editing events in a lineage given the normalized final editing rate: 1 – (10x)^(1/generations), which we validated with simulations. Each editing rate was then passed to Counterfeiter for all sites as a parameter for the editing simulations. Both simulations were run with varying drop rates, where recorders were simulated to the final generation, and individual recorder integrations were dropped randomly to the given proportion. Trees were assembled with IQ-Tree^42^, and analysis and downstream processing were performed with R and TreeUtils (https://github.com/mckennalab/TreeUtils). Normalized Robinson-Foulds comparisons between simulated trees and ground truth were performed using the RF.dist function from the phangorn^57^ package.

Each population was aggregated for the in vitro cell lineage data, and the mean editing rate per site was calculated across all recorders. This was used to create a correlation matrix between samples, and a lineage tree across samples was created using hierarchical clustering. The closest neighbor for each tip was then calculated using the cophenetic.phylo method in the Ape^58^ R package.

## Acknowledgments

We would like to greatly acknowledge the support of Gail Coleman, whose gift directly funded this work. We thank the members of the McKenna lab for experimental help and scientific input. We recognize the Dartmouth Cancer Center and the shared resources supported by the NCI Cancer Center Support Grant 5P30CA023108. We especially thank Fred W. Kolling IV, Laurent Perreard, Carol Ringelberg, and Elizabeth Sergison of the GMBSR for their sequencing guidance and expertise. All Illumina sequencing was carried out in the Genomics and Molecular Biology Shared Resource (RRID:SCR_021293), which is additionally supported by NIH S10 (1S10OD030242) awards. Single-cell studies were conducted through the Dartmouth Center for Quantitative Biology in collaboration with the GMBSR with support from NIGMS COBRE (P20GM130454) and NIH S10 (S10OD025235) awards. This work was supported by N.I.H. award DP2GM149750 and the V Foundation, and A.M. is supported by The Pew Biomedical Scholars program.

## Supplemental Figures

**Supplement Figure 1.**
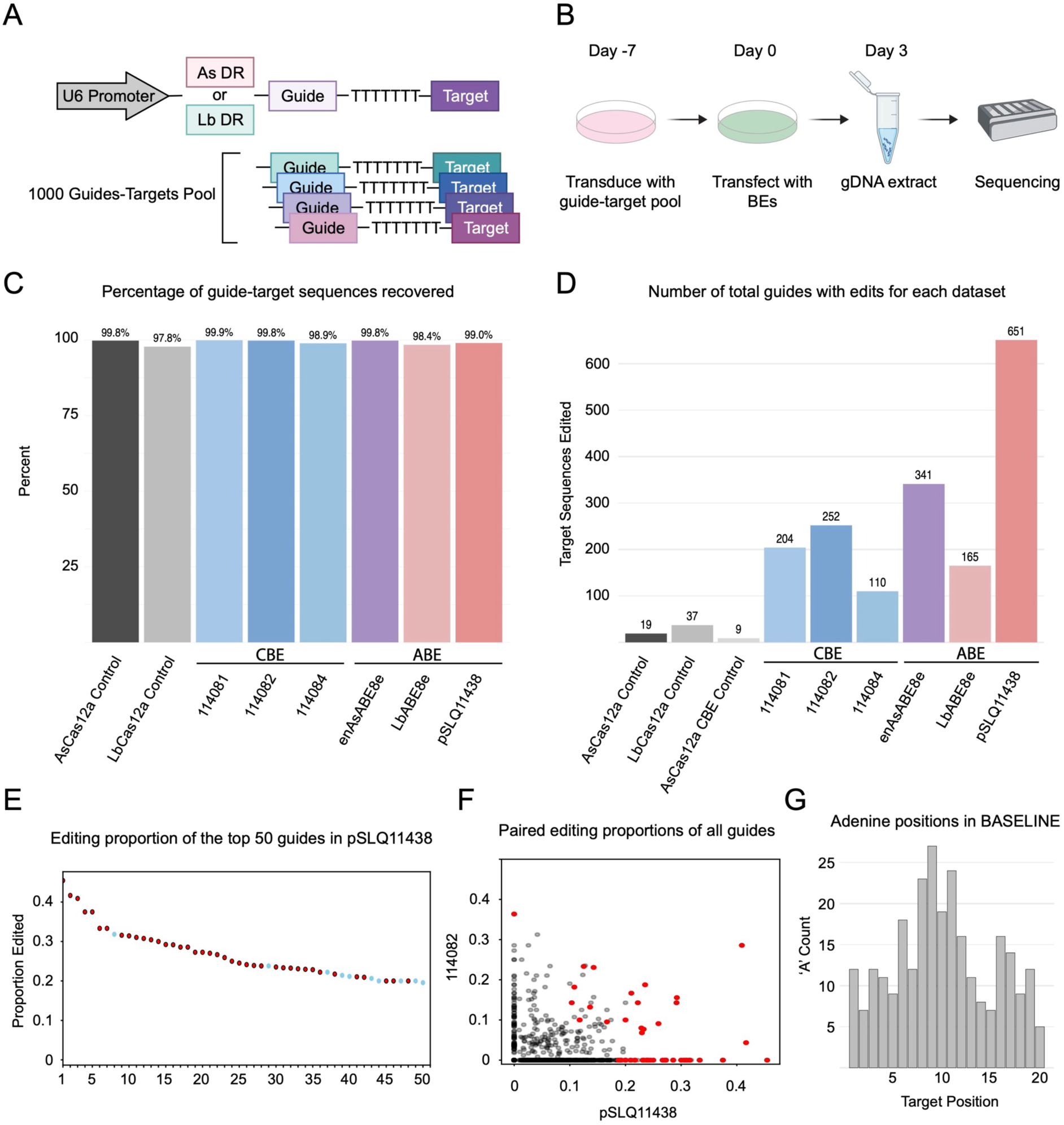
Selecting BASELINE target sequences and Cas12a nucleases. **A)** Schematic for design for cloning of the oligo pool used to test 1000 target sequences. **B)** Experimental workflow for testing 1000 target sequences. **C)** Bar plot that shows the target-guide recovery for each of the tested Cas12a nucleases and no-Cas controls. **D)** Bar plot that shows the number of edited target sequences for each Cas12a base editor sample. **E)** Dot plot that shows the top 50 most highly edited target sequences in the pSLQ11438 sample. Red dots indicate target sequences that were used in the final version of BASELINE. **F)** Dot plot showing the pSLQ11438 (x-ais) and 114082 (y-axis) editing rates for all target sequences. Red dots indicate target sequences that were used in BASELINE. **G)** Histogram that shows sum of adenines across the target positions for all 50 BASELINE targets. Selected targets contained the majority of adenine bases between position 8 to 12.

**Supplemental Figure 2.**
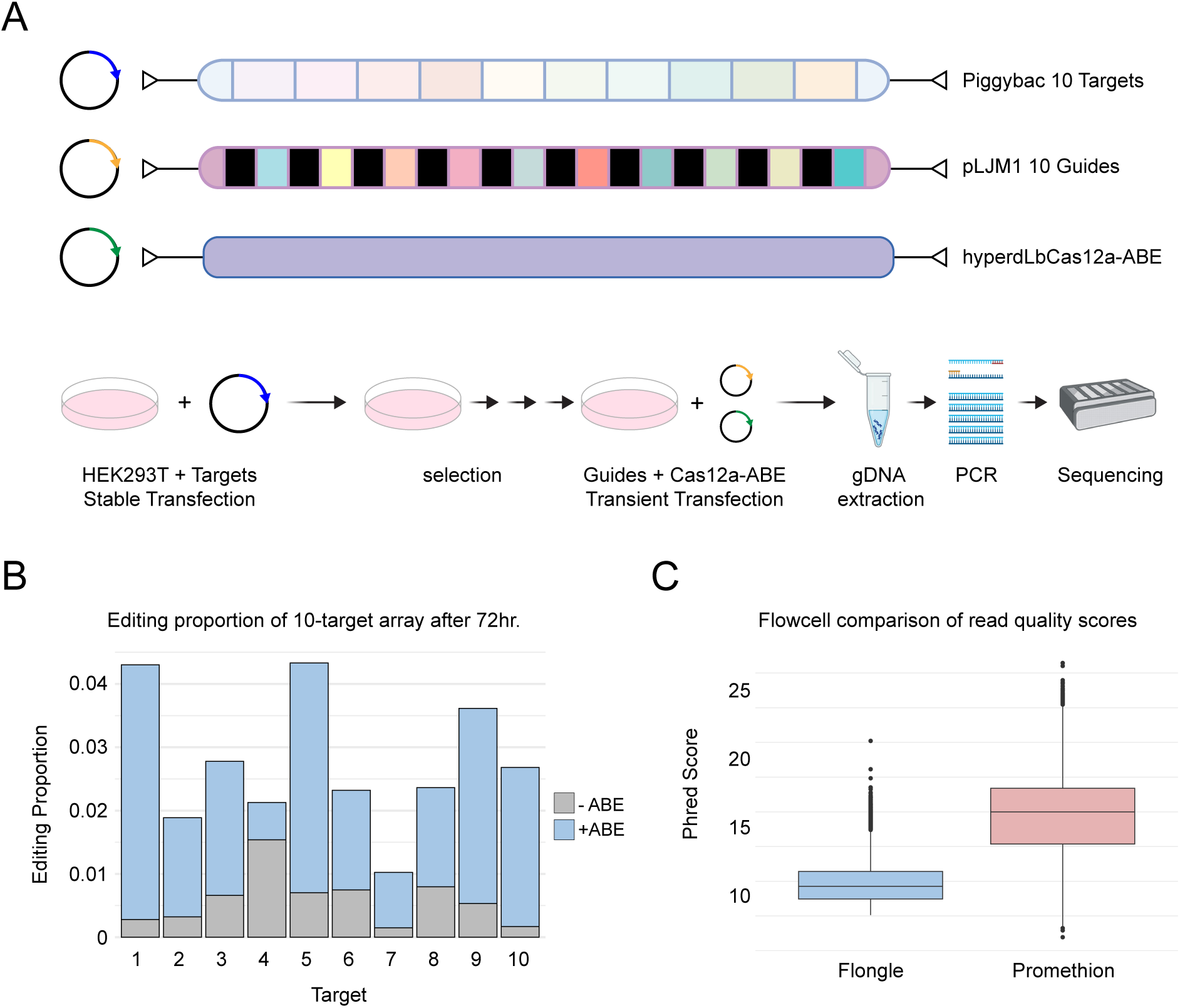
Testing a 10-target multiplexed guide and target recorder. **A)** Schematic for the construct design and experimental workflow for the 10-guide and 10-target multiplex experiment. **B)** Overlapping bar plot shows the edit proportion of the 10-target recorders with and without Cas12a-ABE (grey = no Cas12a-ABE, blue = with Cas12a-ABE). Calculated edit proportion may also include background nanopore sequencing error. **C)** Boxplot that shows the raw read quality scores between the Oxford Nanopore Flongle R10.4.1 (blue) and PromethION R10.4.1 (red) flow cells.

**Supplemental Figure 3.**
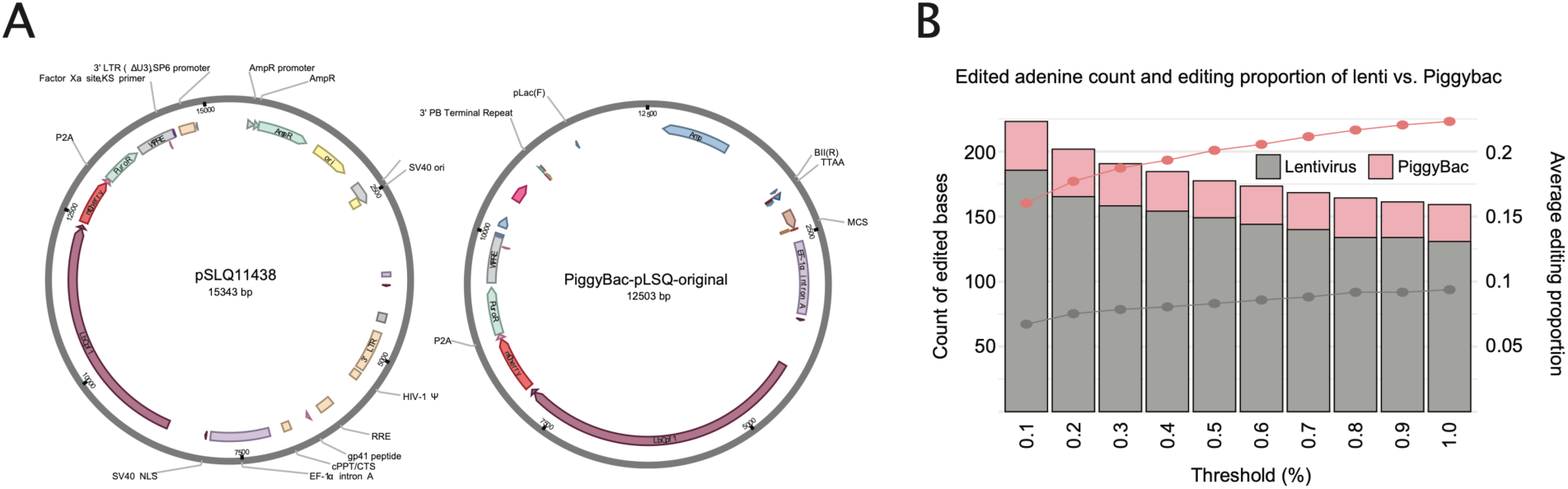
Comparing PiggyBac and lentiviral base editing rates demonstrate that piggyBac accumulates more edits at a higher rate. **A)** Plasmid maps of pSLQ11438 (lentiviral) and PiggyBac-pLSQ-original (piggyBac). **B)** Bar plot and line graph show total edited bases and average edit proportion across 10 different editing thresholds.

**Supplemental Figure 4.**
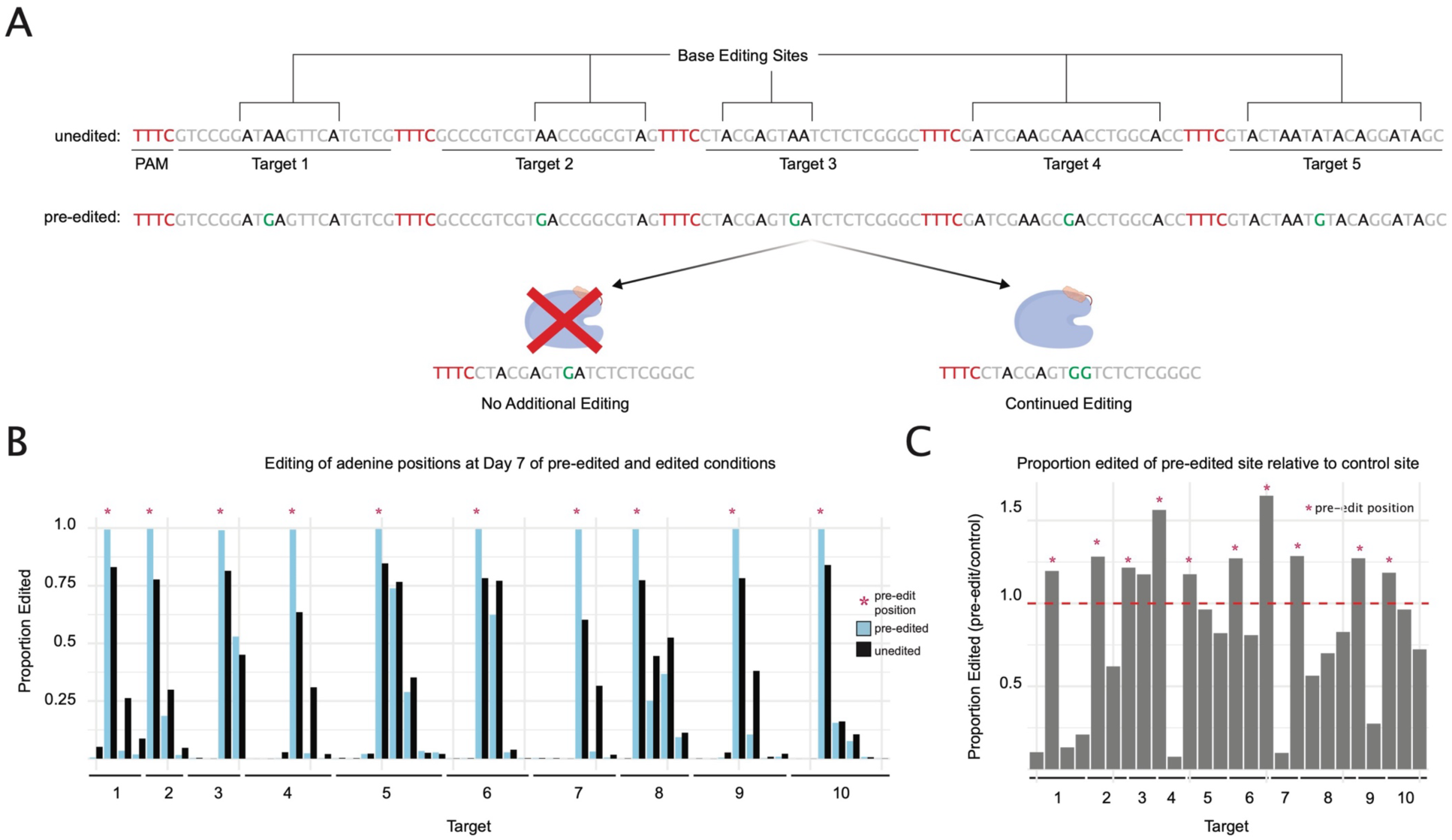
Guide-Target mismatches marginally disrupt successive editing. **A)** Schematic showing an unedited and pre-edited target array and how those A-to-G mutations (in the form of pre-existing edits) count could alter the successive editing potential at a target site. **B)** Bar plot of the editing proportion of all adenine bases in the first 10 targets with and without target pre-editing. Blue bars are from the pre-edited sample. Black bars are from the unedited sample. Red asterisk indicates the site that was pre-edited. **C)** Bar plot detailing the ratio of edit proportion in the pre-edited sample relative to the unedited sample. Bars above the red dashed line indicate more editing in the pre-edited sample. Bars below the red dashed line indicate more editing in the unedited sample. Target sites filtered for adenines that showed more than 0.05 proportion edited in either the unedited or pre-edited sample. Red asterisk indicates the site that was pre-edited.

**Supplemental Figure 5.**
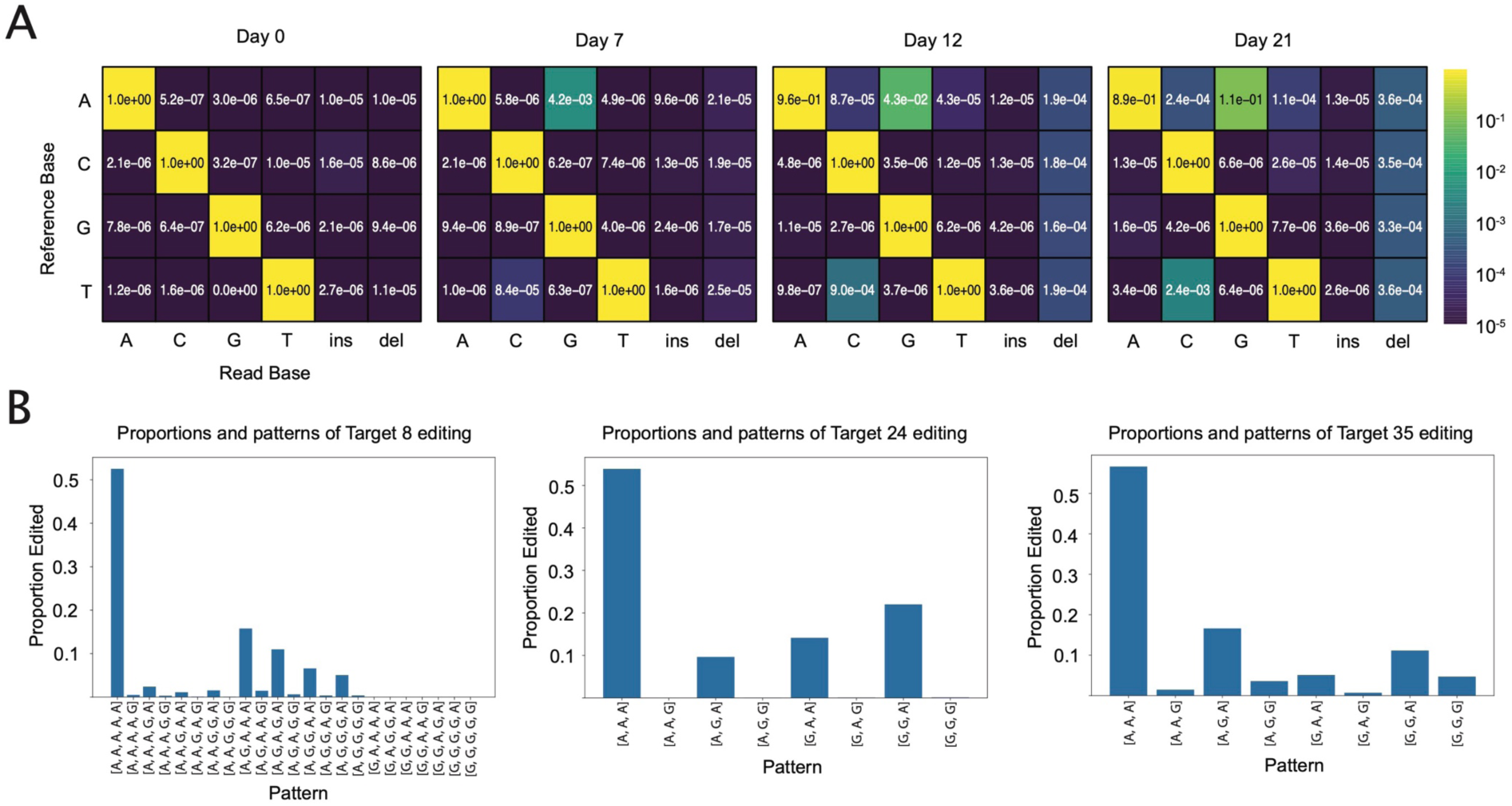
Canonical editing accumulates over time and uniquely across BASELINE targets. **A)** Heatmap conversion matrices of timecourse sequencing data show low off-target error rates with appreciable A-to-G editing. By day 12 and day 21 there is also appreciable T-to-C editing, indicating editing of the non-target strand. **B)** Representative editing outcomes from targets 8, 24, and 35 show variability in editing patterns, regardless of the adenine positions per target.

**Supplementary Figure 6.**
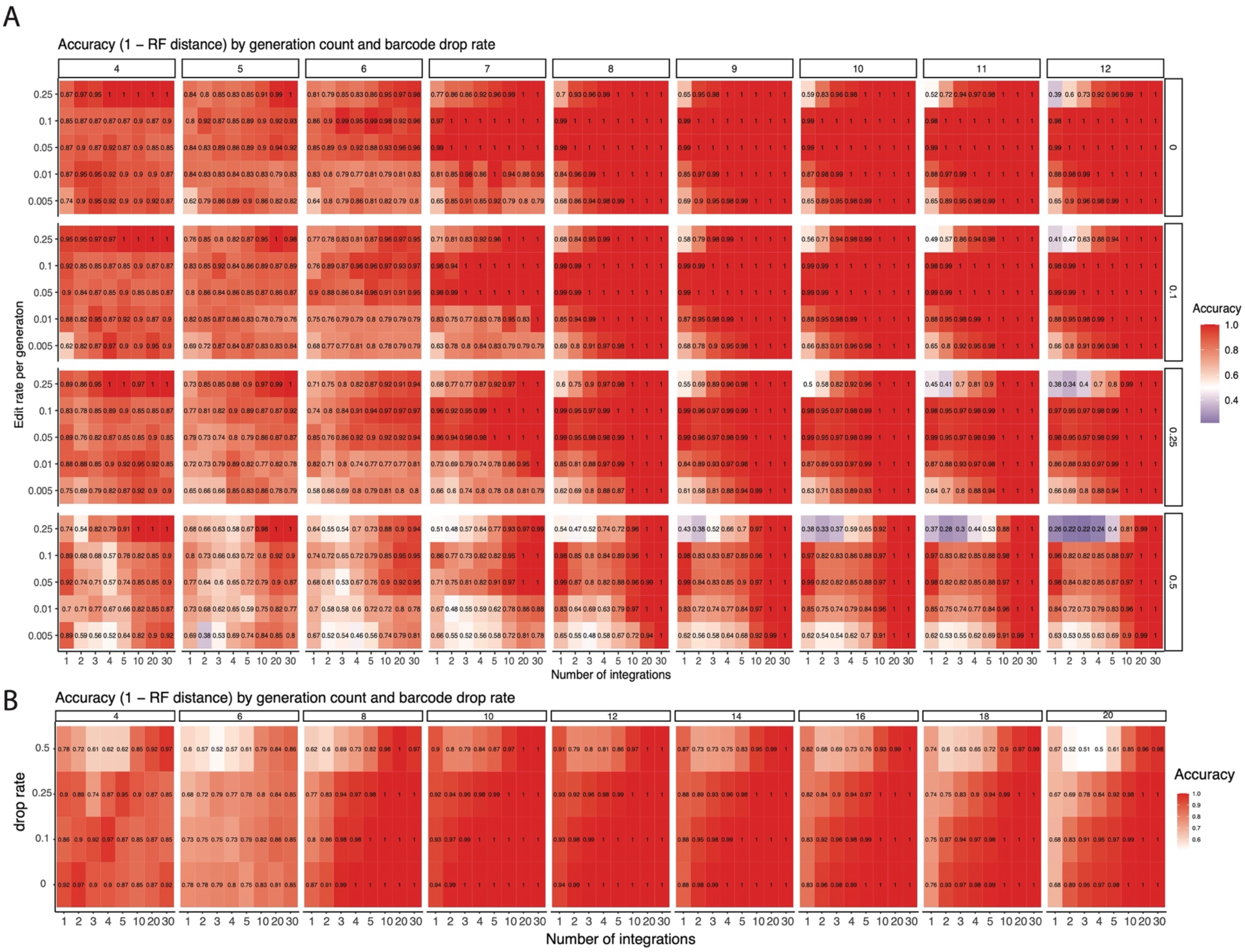
Simulated Lineage tree accuracy. **A)** We simulated lineage trees with various number of 50-target recorder constructs (1-30, x-axis), various editing rates per-cell-generation (y axis), overall tree size (columns), and recorder drop rates (left axis). B) Simulations as in A, with editing rates established from the in vitro timecourse from 4 to 20 generations (∼1 million cells).

**Supplemental Figure 7.**
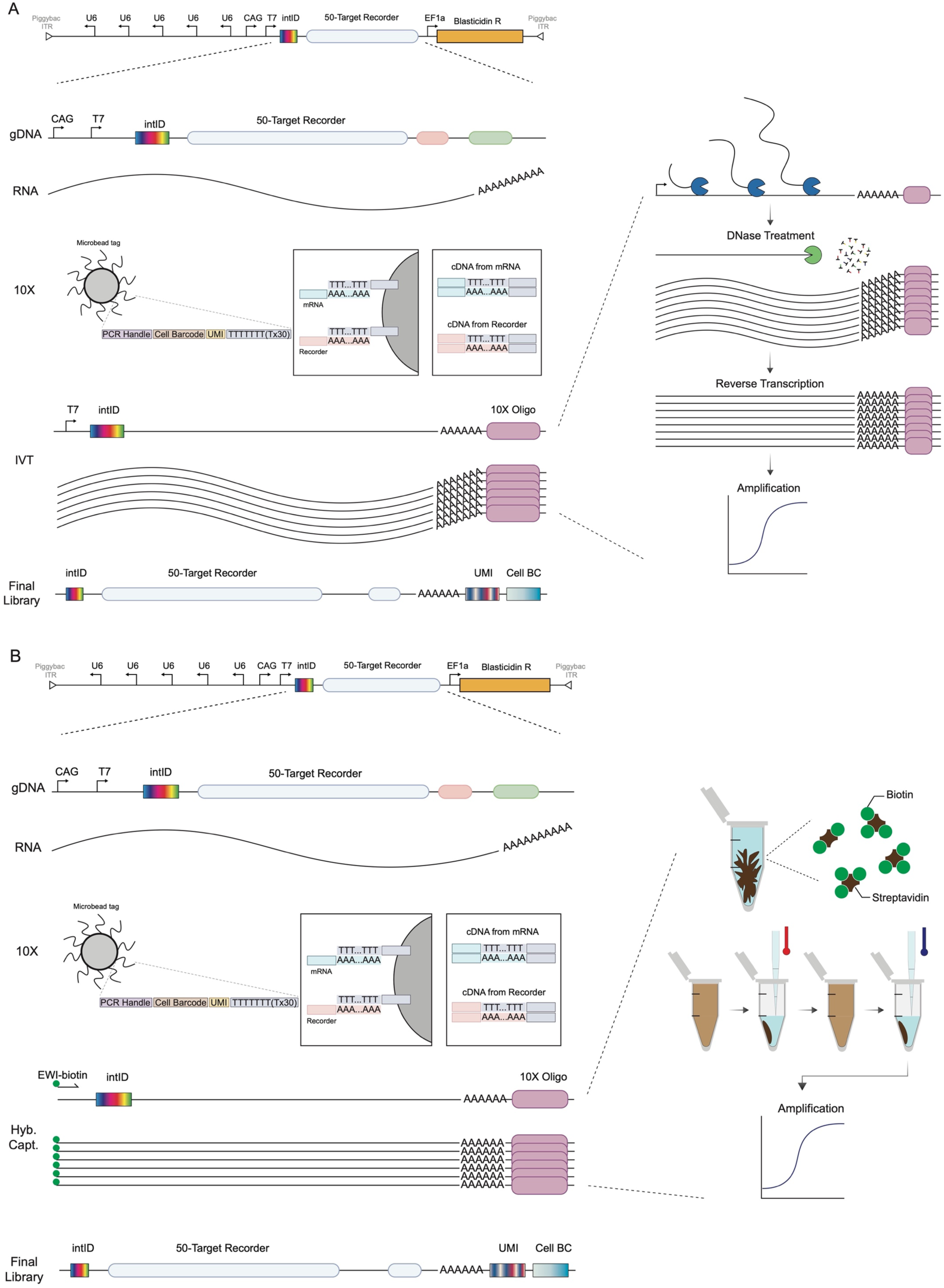
Single-cell sequencing BASELINE schematic. scRNAseq library prep workflow with **A)** IVT and with **B)** hybridization capture. Our final BASELINE construct design contained a T7 promoter sequence and drop-in restriction site for a degenerate integration identifying sequence (intID). By including this static tag at the 5’ end of the recorder we would be able to demultiplex cells by their intIDs before reconstructing phylogenetic trees. Prior to sequencing we generated clean amplicon products from both capture oligos.

**Supplemental Figure 8.**
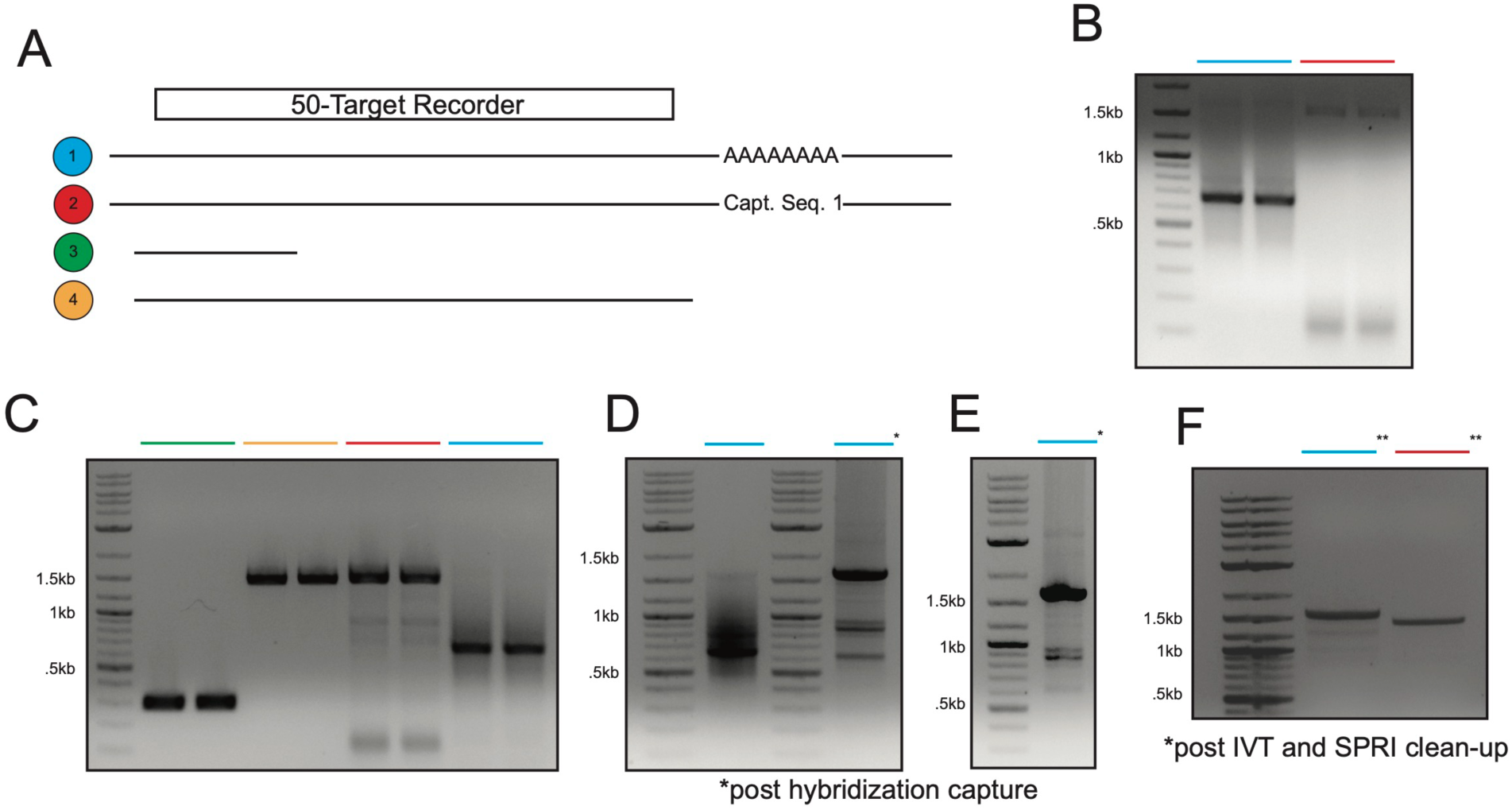
BASELINE recorder isolation and purification by hybridization capture and IVT. **A)** Diagram for PCRs shown in the following panels. Primers EWI & FDZ = Blue, EWI & EWE = Red, EFV & EFT = Green, EFV & ELV = Orange. **B)** - **H)** Agarose DNA gels showing the isolation and purification of BASELINE recorders. **B)** and **C)** PCR using template cDNA from 10X Chromium v4 scRNAseq protocol. **D)** Recorder PCR before and after hybridization capture. **E)** Recorder PCR after hybridization capture, more starting material. **F)** Recorder PCR after T7 IVT and RT.

**Supplemental Figure 9.**
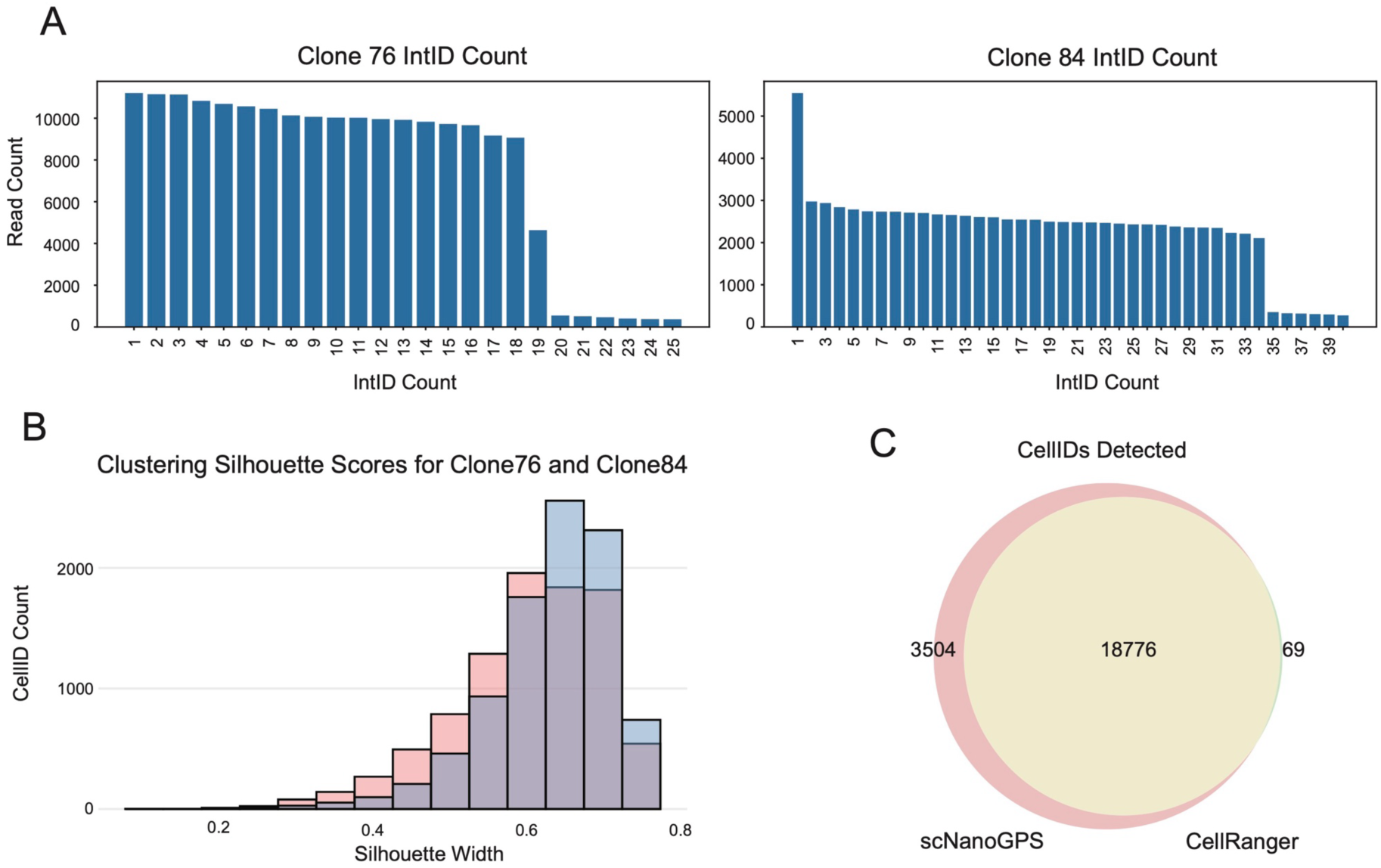
Characterization, capture, and clustering of clone 76 and 84. **A)** Histogram to show the BASELINE recorder integration (intID) diversity of clone 76 (left) and clone 84 (right). **B)** Silhouette scores for hierarchical clustering method (blue = clone 76, red = clone 84). **C)** Venn diagram shows the overlap of detected cellIDs from scNanoGPS and CellRanger.

**Supplemental Figure 10.**
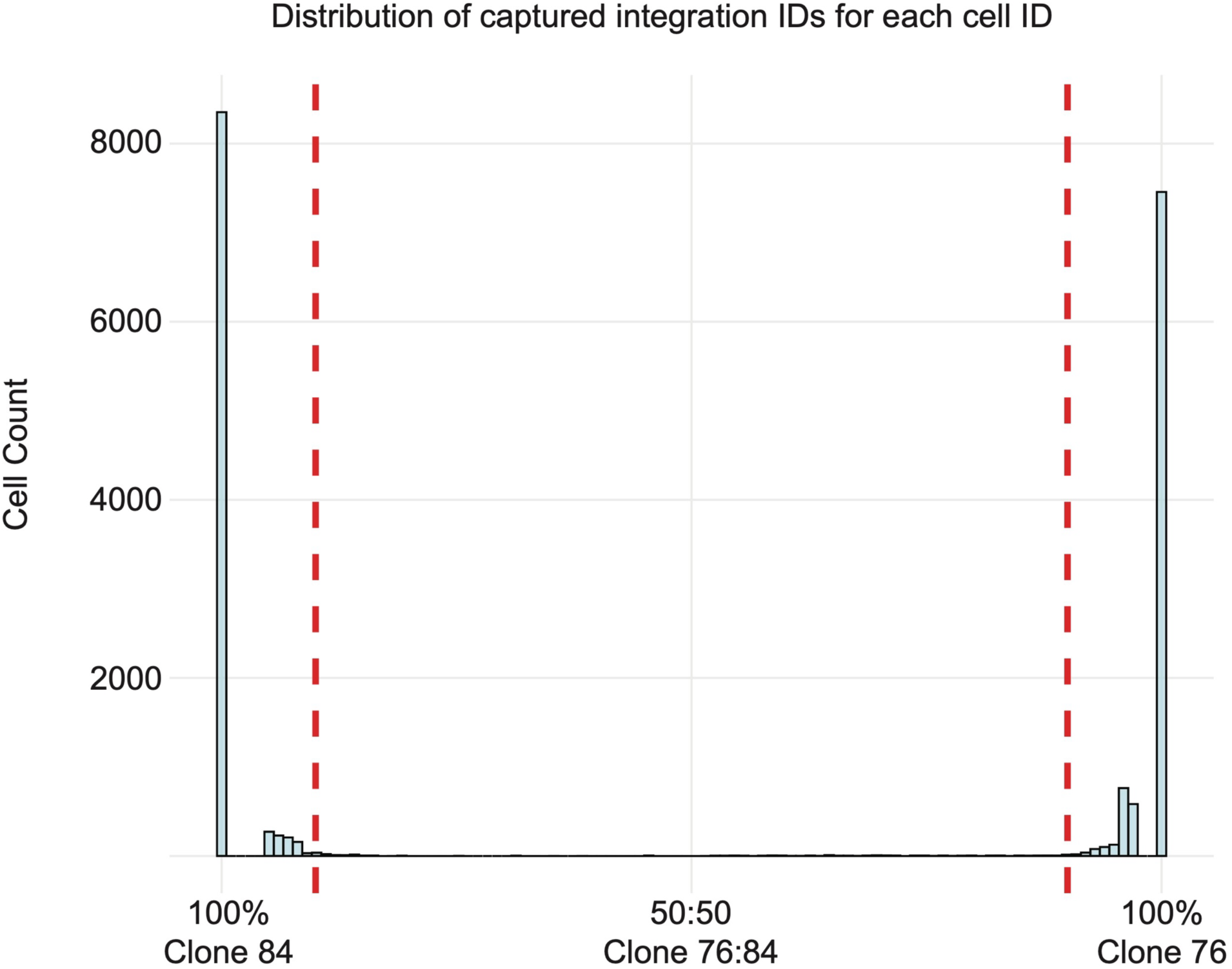
Captured recorder clonal identity per cell. Histogram showing the distribution of intIDs recovered from 18,776 cells. Left-shifted bars indicate cells that have more intIDs belonging to clone 84. Bars on right indicate cells that have more intIDs belonging to clone 76. Vertical red dashed lines indicate 90% cut-off for captured intIDs per cell.

**Supplemental Figure 11.**
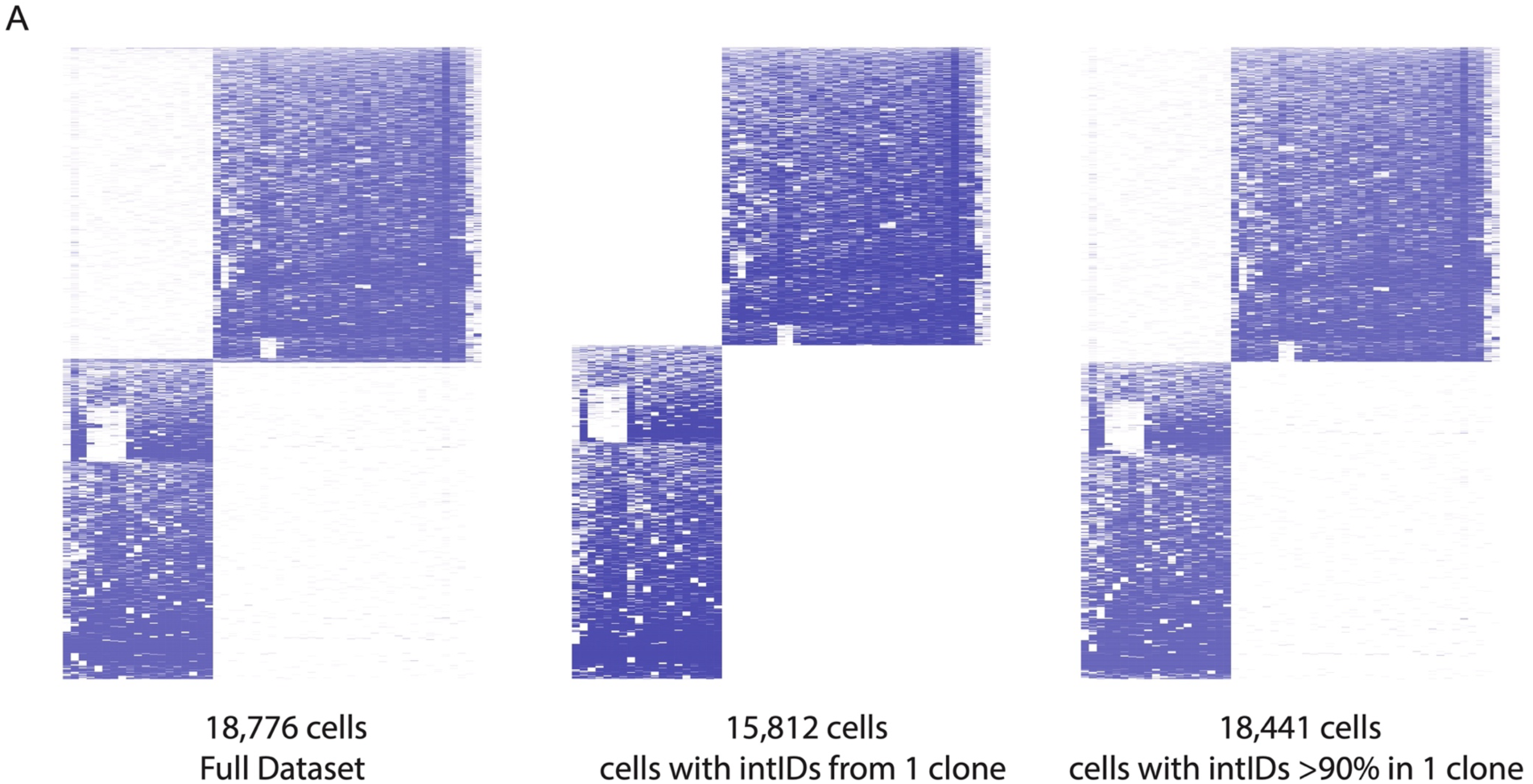
Heirarchical clustering of cellIDs and intIDs recovered from scNanoGPS. Clustering of all 18,776 cells (left). Clustering of only cells that contained purely intIDs from either clone 76 or clone 84 (middle). Clustering of cells that contained greater than or equal to 90% intIDs from either clone 76 or clone 84 (right).

**Supplemental Figure 12.**
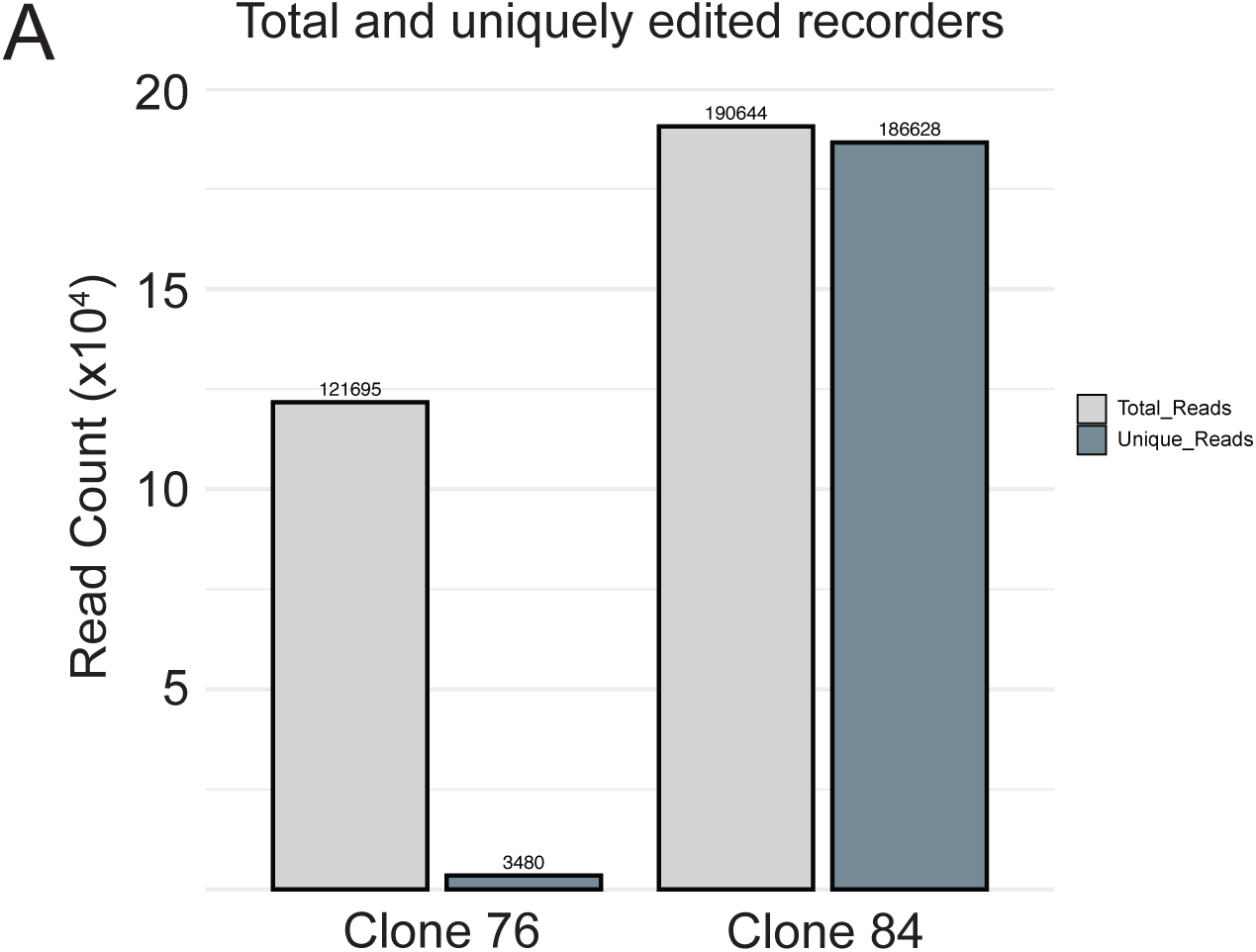
Edited recorders are more unique in clone 84 than clone 76. **A)** Total edited reads without ambiguous bases for clone 76 and clone 84 populations show that clone 84 recorders remain largely unique compared to clone 76.

**Supplemental Figure 13.**
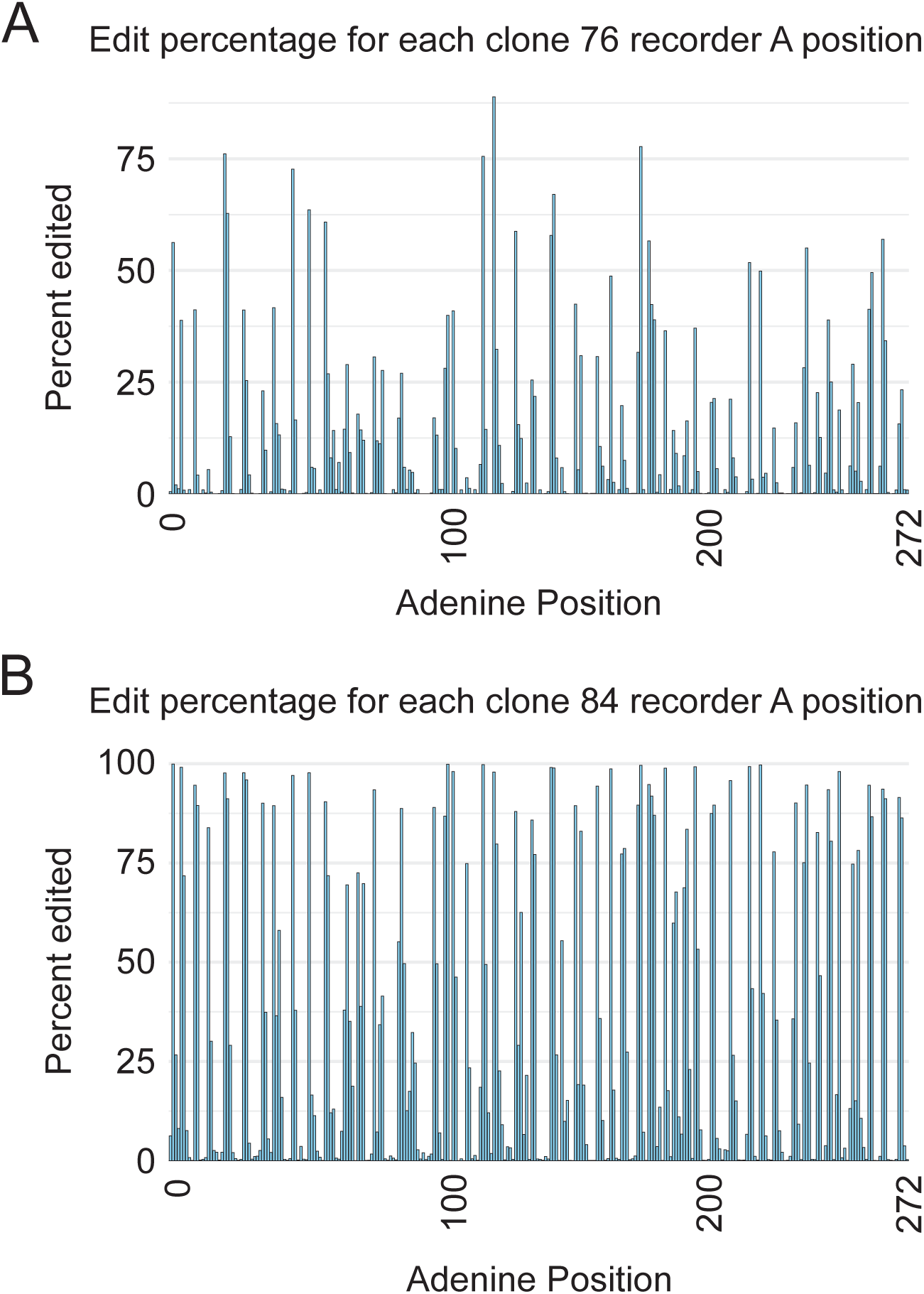
The calculated edit percentage for all BASELINE adenine positions shows saturated events in clone 84. **A) and B)** Bar plots show the edit percentage for all adenine positions in BASELINE. No adenines have been edited across all captured recorders in clone 76 while XX adenine positions are saturated (>99%) with edits in clone 84.

**Supplemental Figure 14.**
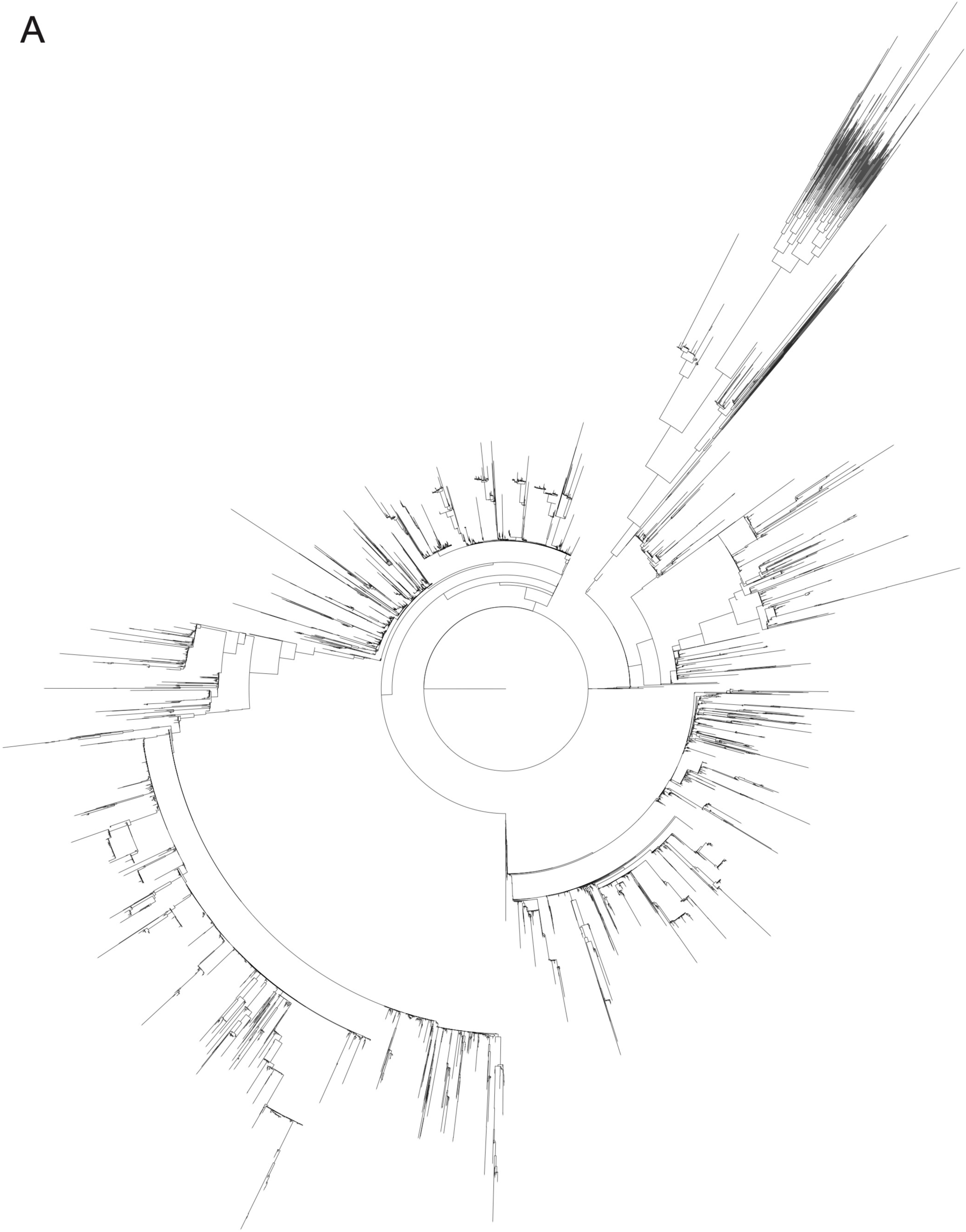
Clone 76 reconstructed phylogenetic tree. Clone 76 shows 9,260 cells reconstructed into a phylogenetic tree.

**Supplemental Figure 15.**
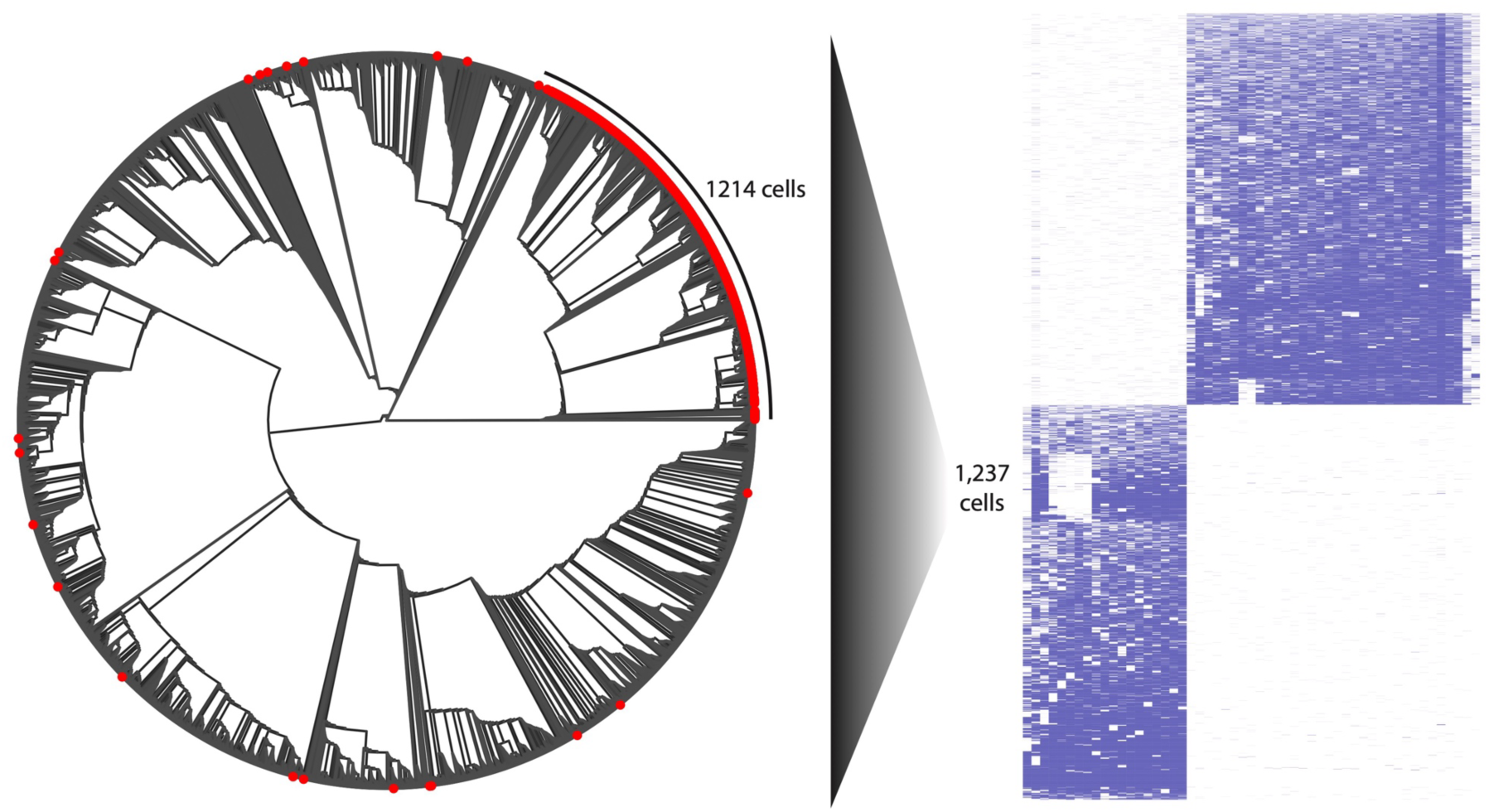
Hierarchical clustered clone 76 cells map together in phylogenetic tree. Phylogenetic tree of clone 76 (left) with hierarchical clustering of full lineage tracing dataset (18,441 cells, right) shows over 98% of cells missing all recorders 1, 4, 5, 6, 7, and 8 map to the same clade of the phylogenetic tree.

**Supplemental Figure 16.**
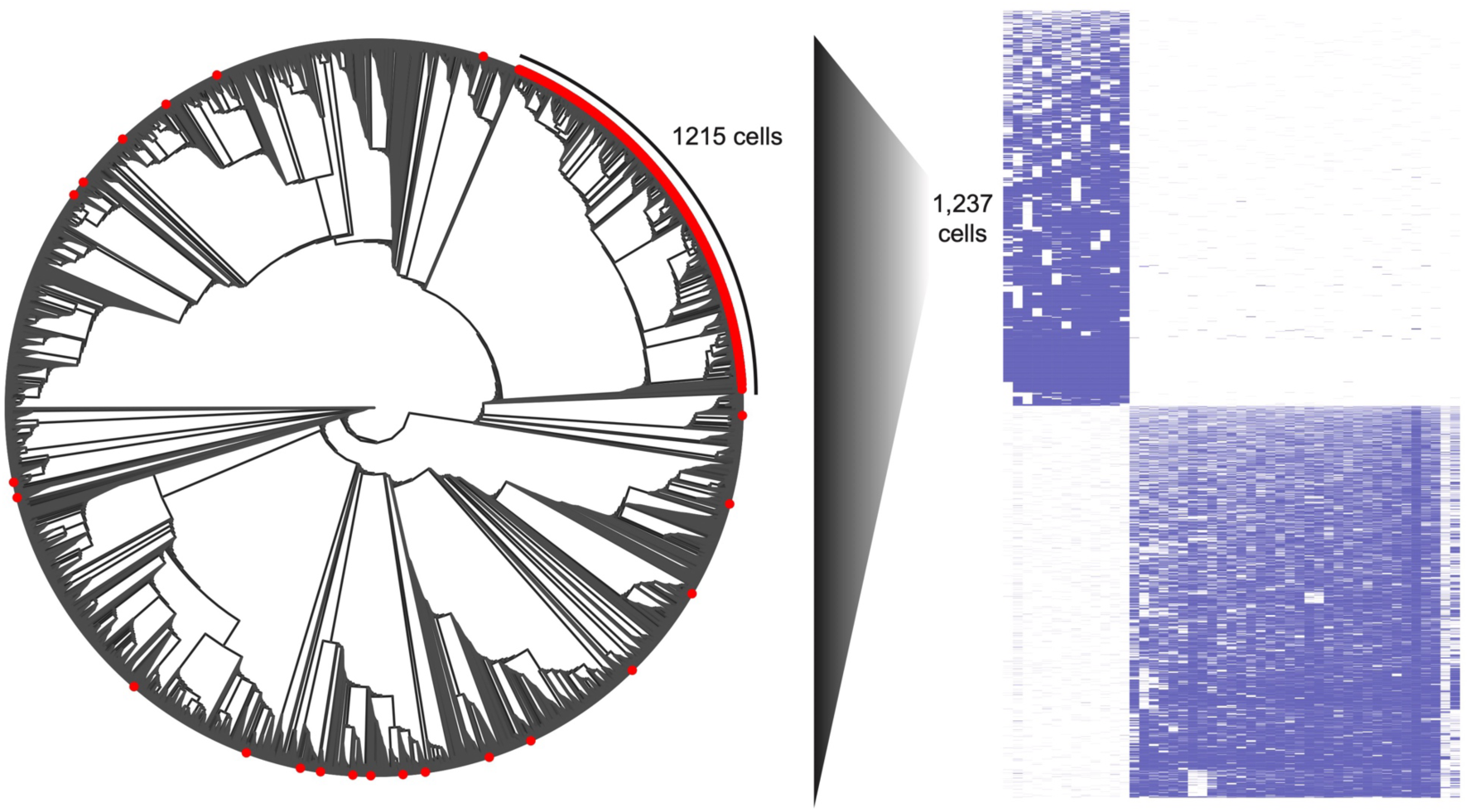
Hierarchical clustering of clone 76 cells after removal of six recorders mapping onto phylogenetic tree. Phylogenetic tree of clone 76 (left) and hierarchical clustering of lineage dataset (right) with removal of six recorders and their lineage information shows that over 98% of the cells that were missing the six recorders map to the same clade of the phylogenetic tree.

